# TXNDC5 Governs Extracellular Matrix Homeostasis in Pulmonary Hypertension

**DOI:** 10.1101/2025.11.21.689856

**Authors:** Zhen Chen, Fanhao Kong, Lixin Huang, Wenze Wu, Ziping Wang, Hanbin Chen, Junting Zhang, Lin Deng, Chengrui Cao, Xiaoyan Zhang, Zixin Liu, Kai-Chien Yang, Wei-Ting Chang, Jin-Song Bian, Xiaowei Nie

## Abstract

**BACKGROUND:** Pulmonary hypertension (PH) is characterized by vascular remodeling without effective treatments. Thioredoxin domain containing 5 (TXNDC5), a member of the protein disulfide isomerases (PDI) family, regulates protein folding and vascular homeostasis, yet its role in PH remains unknown.

**METHODS:** Label-free proteomics profiled protein expression in lungs from PH patients. TXNDC5 was analyzed by single-cell RNA sequencing, immunofluorescence, and Western blot. Endothelial gain- and loss-of-function approaches were applied in Sugen5416/hypoxia (SuHx)-induced rodent PH models. RNA sequencing and protein-protein interaction analysis were used to investigate underlying mechanisms.

**RESULTS:** TXNDC5 was significantly upregulated in the lungs of patients with PH and in experimental PH models, with predominant localization in endothelial cells (ECs) of remodeled distal pulmonary arteries. Endothelial TXNDC5 overexpression exacerbated pulmonary vascular remodeling, elevated right ventricular systolic pressure, and promoted right ventricular hypertrophy, whereas global or endothelial-specific TXNDC5 deficiency conferred protection against SuHx-induced PH. Hypoxia-induced factor (HIF)-2α transcriptionally activated TXNDC5 to drive PH development. Single-cell RNA sequencing identified a distinct subpopulation characterized by TXNDC5^high^ extracellular matrix (ECM)-producing ECs. Bulk RNA sequencing combined with protein-protein interaction analysis revealed that TXNDC5 regulated ECM homeostasis through biglycan (BGN). Pharmacological inhibition of TXNDC5 with E64FC26 and endothelial-targeted TXNDC5 gene therapy significantly attenuated PH severity in rats.

**CONCLUSIONS:** Our study reveals that TXNDC5 is a main modulator to regulate ECM homeostasis and may serve as a promising target for the treatment of PH.

## INTRODUCTION

Pulmonary hypertension (PH) is a serious cardiovascular disease characterized by elevated pulmonary artery pressure and right heart failure, with high morbidity and mortality^1^. Dysfunction of pulmonary artery endothelial cells (PAECs) under hypoxia initiates vasoconstriction^2^, while pulmonary artery smooth muscle cells (PASMCs) become hyperproliferative and apoptosis-resistant phenotype, leading to vascular remodeling^3^. However, there remains no curative clinical treatment option for reversing remodeled vessels against the disease^4,5^. It highlights the need for further exploration of PH pathogenesis and new therapeutic targets.

Protein disulfide isomerase (PDI) is an endoplasmic reticulum enzyme family that mediates disulfide bond formation, protein folding, and regulates endothelial function to maintain vascular homeostasis^6,7^. Thioredoxin domain containing 5 (TXNDC5), mainly expressed in endothelial cells (ECs) and also called endothelial PDI, is linked to fibrosis^8^, diabetes^9^, rheumatoid arthritis^10^, and cancers^11^. Inhibition of TXNDC5 restores endothelial dysfunction to mitigate vascular calcification and atherosclerosis^12,13^. However, its role in pulmonary vascular remodeling in PH remains unclear.

Extracellular matrix (ECM) is a complex network of proteins and polysaccharides secreted by cells, providing structural and communicative support to adjacent cells^14^. Remodeling of pulmonary arterial ECM occurs in the early stage of PH pathogenesis, before increased wall thickness and pulmonary artery pressure. ECM remodeling is the cause rather than a consequence of pulmonary vascular remodeling in PH^15^. Recently, we revealed that the ECM proteins in PAECs accelerated pulmonary vascular remodeling through interaction with hypoxia-induced factor (HIF)-1α in a positive feedback loop manner, providing evidence for early intervention in ECM remodeling to prevent PH progression^16,17^.

In this study, we identified TXNDC5 as a markedly dysregulated factor in ECs within the pulmonary vasculature in PH. Global or endothelial-specific deficiency of TXNDC5 alleviated Sugen5416/hypoxia (SuHx)-induced PH, whereas TXNDC5 overexpression aggravated PH. HIF-2α was identified as a transcriptional activator of TXNDC5. Single-cell RNA sequencing identified a distinct subpopulation characterized by TXNDC5^high^ ECM-producing ECs. TXNDC5 directly interacts with biglycan (BGN) and facilitates its folding, thereby promoting pathological ECM remodeling. Pharmacological inhibition or pulmonary endothelium-targeted silencing of TXNDC5 effectively mitigated PH in rats. The present study demonstrated that TXNDC5 may be a key protein to control ECM homeostasis and a promising therapeutic target for PH.

## METHODS

Expanded methods and materials are available in **Supplemental Material 1**.

### Data Availability Statement

Source data and original blots or gel images for this study are available from the corresponding author upon reasonable request.

## RESULTS

### TXNDC5 Expression is Increased in PH Patients and Rodent Models

We performed 4D label-free proteomic profiling of lung tissues from 5 patients with PH and 5 non-PH donors (**Figure 1A** and **Supplemental Material 2**). Among the human PDI family proteins, TXNDC5 was the most markedly dysregulated protein in PH patients compared with donor controls (fold change=3.45, *P*=0.0048) (**Figure 1B**). Consistently, increased TXNDC5 protein levels in lungs from PH patients were confirmed (**Figure 1C**), and similar results were recapitulated in multiple rat and mouse PH models (**Figure 1D** and **Figure S1A-B**).

**Figure 1.**
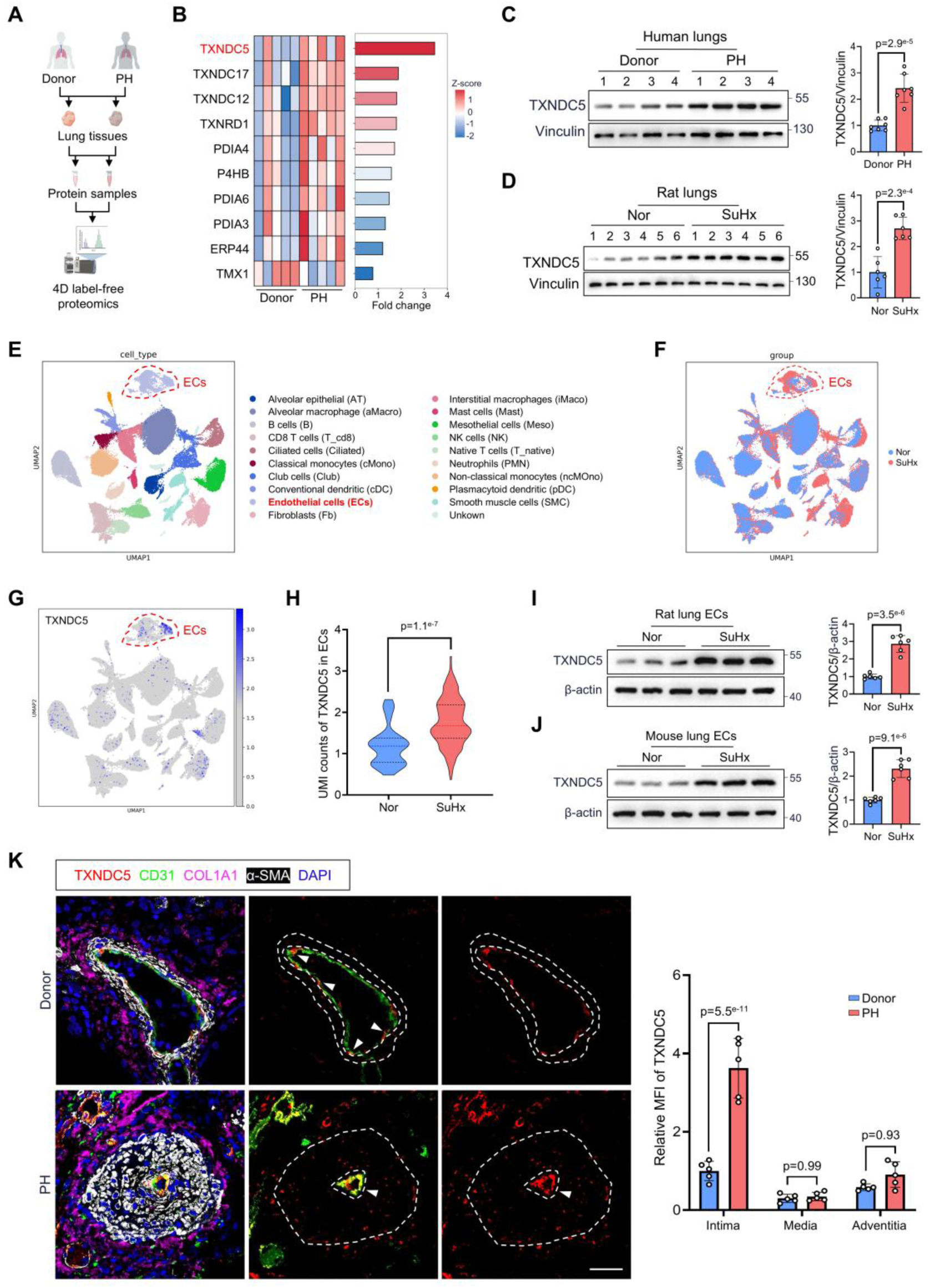
TXNDC5 was upregulated in PH. **A**, Schematic illustration showing the 4D label-free proteomics of lungs from non-PH donors and patients with PH. **B**, Heatmap presenting differential expression of PDI proteins, highlighting TXNDC5 among the members (n=5). **C**, Western blots showing TXNDC5 protein levels in lungs from donors and PH patients (n=7). **D**, Western blots showing TXNDC5 protein levels in lungs from SuHx-treated rats and normoxic controls (Nor) (n=6). **E,** UMAP visualization displaying annotated lung cell types, with endothelial cells (ECs) highlighted. **F,** UMAP plots showing the distribution of lung cell populations from Nor and SuHx-exposed rats. **G**, Feature plot showing TXNDC5 expression across lung cell populations. **H**, Violin plot demonstrating TXNDC5 expression in ECs quantified by unique molecular identifier (UMI) counts. **I** and **J**, Western blots showing TXNDC5 expression in primary lung ECs from SuHx-treated PH rats and mice compared with controls. **K**, Multiplex immunofluorescence staining showing TXNDC5 (red), CD31 (green), α-SMA (white), and COL1A1 (purple) in pulmonary arteries from human lungs (n=5). Nuclei were counterstained with DAPI (blue). Scale bar=25 μm. MFI indicates mean fluorescence intensity. Data represent the mean±SD. Comparisons of parameters were performed with the unpaired 2-tailed Student *t*-test (**C-D** and **I-J**) or 2-tailed Mann-Whitney *U* test (**H**) or 2-way ANOVA (**K**), followed by Tukey honestly significant difference test for multiple comparisons. PDI indicates protein disulfide isomerase; α-SMA, α-smooth muscle actin; COL1A1, collagen type I alpha 1; DAPI, 4ʹ,6-diamidino-2-phenylindole; Nor, normoxia; and SuHx, Sugen5416/hypoxia.

To examine the cellular localization of TXNDC5 in PH, we analyzed a single-cell RNA sequencing dataset (GSE273062) from lung tissues of SuHx-exposed rats and normoxic controls (Nor) (**Figure 1E-F**). TXNDC5 was primarily expressed in ECs (**Figure 1G**) and was markedly upregulated in ECs from SuHx lungs compared with controls (**Figure 1H**). Western blot analysis of primary lung ECs corroborated these findings (**Figure 1I-J**). In pulmonary arteries from human lungs, multiplex immunofluorescence revealed strong co-localization of TXNDC5 with the endothelial marker CD31. Moreover, TXNDC5 expression was markedly increased in the intimal layer of remodeled pulmonary arteries from PH patients (**Figure 1K**). Taken together, these findings suggest that the TXNDC5 may be a critical dysregulated factor in ECs within the pulmonary vasculature that contributes to PH pathogenesis.

### TXNDC5 Mediates the Development of SuHx-Induced PH

To investigate the role of TXNDC5 *in vivo* in the development of experimental PH, TXNDC5 knockout (TXNDC5^-/-^) mice were challenged with or without SuHx (**Figure 2A**). TXNDC5 deletion alone had no significant effects but markedly reduced RVSP (**Figure 2B**) and Fulton index (right ventricle weight to left ventricle plus interventricular septum weight, ie, RV/(LV+S)) (**Figure S1C**) induced by SuHx. Echocardiography indicated improved right ventricular (RV) function, with increased pulmonary artery acceleration time/ pulmonary artery ejection time ratio (PAT/PET), tricuspid annular plane systolic excursion (TAPSE), and right ventricular fractional area change (RVFAC), along with reduced right ventricular free wall thickness (RVFWT) (**Figure S1D**). In addition, H&E and α-SMA staining demonstrated attenuation of distal pulmonary artery medial wall thickening and muscularization under SuHx (**Figure 2C-D**). Consistent with the pulmonary arterial changes, wheat germ agglutinin (WGA) staining proved that the RV cardiomyocyte hypertrophy was markedly inhibited in SuHx-treated TXNDC5^-/-^ mice when compared with their control littermates (**Figure 2E**).

**Figure 2.**
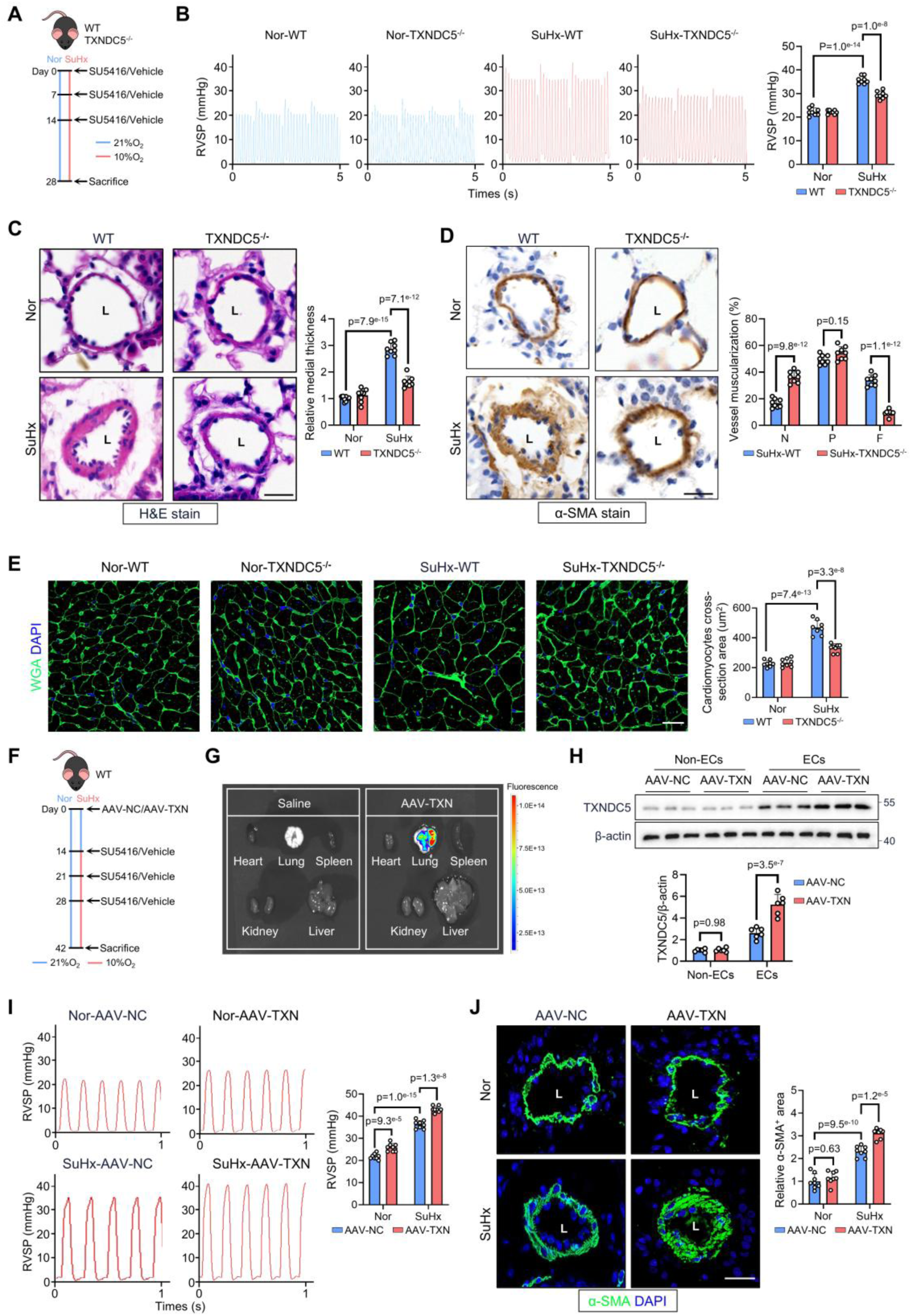
TXNDC5 contributed to SuHx-induced pulmonary vascular remodeling. **A**, Schematic overview of the SuHx-induced PH protocol in wild-type (WT) or TXNDC5 global knockout (TXNDC5^-/-^) mice. **B**, Hemodynamic measurements showing the tracings and quantification of right ventricular systolic pressure (RVSP) in WT and TXNDC5^-/-^ mice (n=8). **C**, Hematoxylin and eosin (H&E) staining of distal pulmonary arteries in WT and TXNDC5^-/-^ mice (n=8). L indicates lumen. Scale bar=25 μm. **D**, Immunohistochemical staining for α-SMA showing the muscularization of distal pulmonary arteries in WT and TXNDC5^-/-^ mice (n=8). L indicates lumen. Scale bar=25 μm. **E**, WGA (green) staining showing the right ventricular morphology in WT and TXNDC5^-/-^ mice (n=8). Nuclei were counterstained with DAPI (blue). Scale bar=20 μm. **F**, Schematic diagram illustrating pulmonary endothelial-specific TXNDC5 overexpression via AAV delivery in mice. **G**, Representative images of major organs from saline-treated control or AAV-TXN-treated mice, lung-targeted fluorescence detected by the IVIS imaging system. **H**, Western blot showing TXNDC5 expression in lung endothelial cells (ECs) after AAV delivery (n=6). **I**, Hemodynamic measurements showing RVSP in AAV-TXN- or AAV-NC-treated mice under normoxic or SuHx conditions (n=8). **J**, Immunofluorescence staining for α-SMA displaying the pulmonary arterial smooth muscle coverage in AAV-TXN- or AAV-NC-treated mice. Nuclei were counterstained with DAPI (blue). L indicates lumen. Scale bar=25 μm. Data represent the mean±SD. Comparisons of parameters were performed with 2-way ANOVA, followed by Tukey honestly significant difference test for multiple comparisons. DAPI indicates 4ʹ,6-diamidino-2-phenylindole; α-SMA, α-smooth muscle actin; N, non-muscularized; P, partially muscularized; F, fully muscularized; WGA, wheat germ agglutinin; Nor, normoxia; and SuHx, Sugen5416/hypoxia.

With a gain-of-function approach, an adeno-associated virus (AAV) serotype engineered to specifically target lung ECs was delivered via the tail vein to overexpress TXNDC5 specifically (AAV-TXN) (**Figure 2F**). *Ex vivo* imaging confirmed lung targeting (**Figure 2G**), and Western blot verified endothelial-specific overexpression (**Figure 2H**). In response to the SuHx challenge, AAV-TXN mice further increased in both RVSP and RV/(LV+S) (**Figure 2I** and **Figure S1E**). Echocardiography analysis showed reduced PAT/PET, TAPSE RVFAC, and increased RVFWT under SuHx (**Figure S1F**). Consistently, immunofluorescence staining demonstrated an increased α-SMA-positive area in distal pulmonary arteries in SuHx-treated AAV-TXN mice (**Figure 2J**). Conversely, endothelial TXNDC5 overexpression was found to exacerbate SuHx-induced PH.

### Endothelial Cell-Specific Deletion of TXNDC5 Ameliorates SuHx-Induced PH

To examine the role of endothelial TXNDC5 in PH, endothelial-specific TXNDC5 knockout mice (TXNDC5^ECKO^) were generated, validated (**Figure S1G**), and challenged by SuHx (**Figure 3A**). Under SuHx, TXNDC5^ECKO^ mice exhibited reduced distal pulmonary arterial wall thickness and α-SMA positive area (**Figure 3B**). Deletion of endothelial TXNDC5 significantly attenuated SuHx-induced increases in RVSP (**Figure 3C**), RV/(LV+S) (**Figure 3D**), and improved RV function **(Figure S1H**). Lung angiography confirmed better intrapulmonary contrast filling in TXNDC5^ECKO^ mice compared with TXNDC5^f/f^ controls, indicating preserved pulmonary vasculature (**Figure 3E**). In addition, SuHx-induced RV hypertrophy was also markedly attenuated, with decreased RV wall thickness and cardiomyocyte cross-sectional area (**Figure 3F-G**). Together, these findings underscore the crucial role of endothelial TXNDC5 in the development of PH and highlight its potential as a sex-independent therapeutic target.

**Figure 3.**
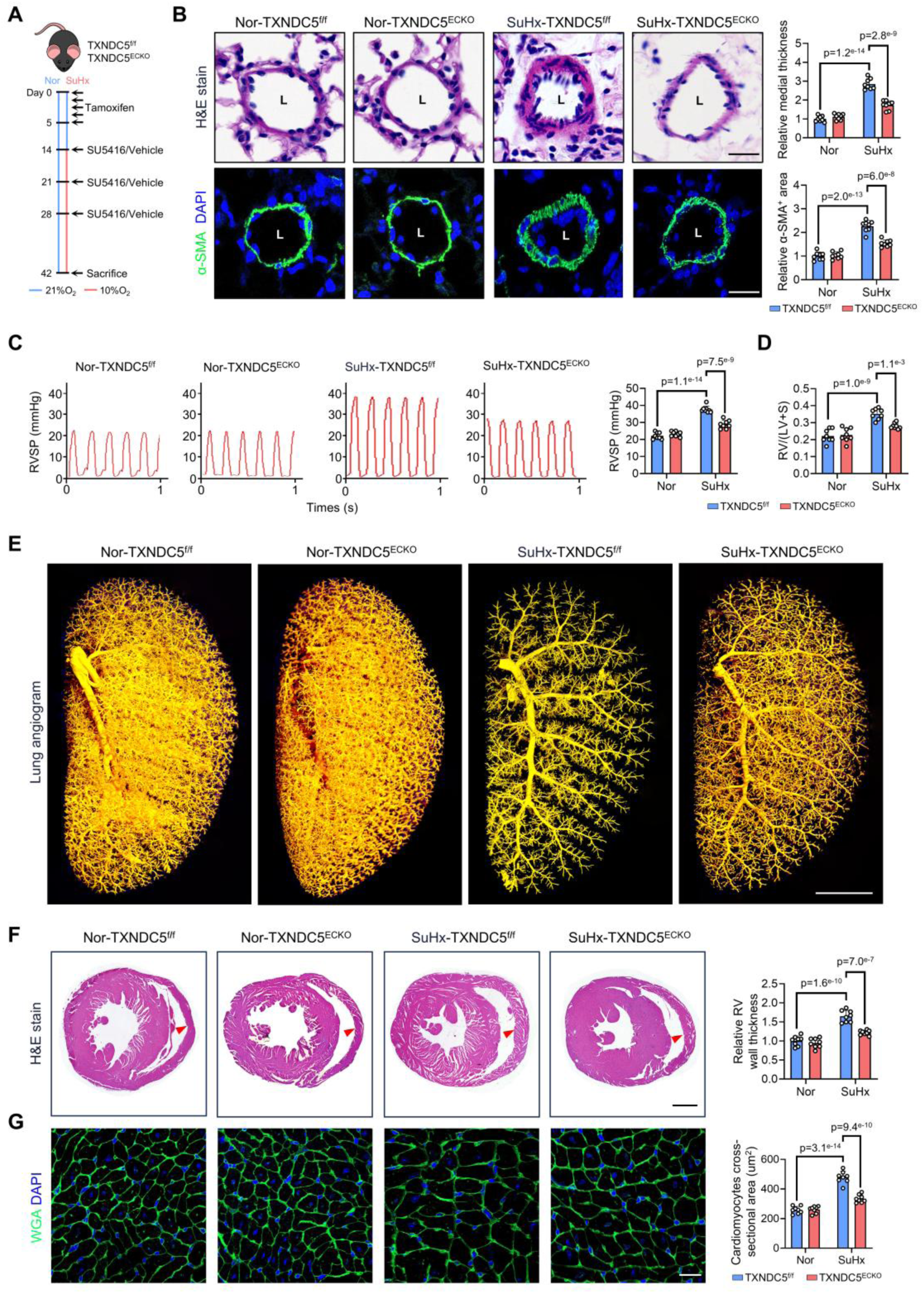
Endothelial-specific deletion of TXNDC5 ameliorated SuHx-induced PH. **A**, Schematic diagram showing the protocol for generating endothelial-specific TXNDC5 knockout (TXNDC5^ECKO^) mice. **B**, Hematoxylin and eosin (H&E) and α-SMA (green) staining showing distal pulmonary arteries in TXNDC5^f/f^ and TXNDC5^ECKO^ mice under the indicated conditions. Nuclei were counterstained with DAPI (blue). L indicates lumen. Scale bar=25 μm. **C**, Hemodynamic measurements showing the right ventricular systolic pressure (RVSP) in TXNDC5^f/f^ and TXNDC5^ECKO^ mice (n=8). **D**, Fulton index [RV/(LV+S)] demonstrating RV hypertrophy in TXNDC5^f/f^ and TXNDC5^ECKO^ mice (n=8). **E**, Lung angiogram of the left lungs showing intrapulmonary contrast filling in TXNDC5^f/f^ and TXNDC5^ECKO^ mice. Scale bar=2 mm. **F**, Histological cross-sections of the mid-right ventricle showing RV wall morphology in TXNDC5^f/f^ and TXNDC5^ECKO^ mice (n=8). Scale bar=1 mm. **G**, Immunofluorescence staining of WGA (green) showing right ventricular cardiomyocyte hypertrophy in TXNDC5^f/f^ and TXNDC5^ECKO^ mice. Nuclei were counterstained with DAPI (blue). Scale bar=20 μm. Data represent the mean±SD. Comparisons of parameters were performed with 2-way ANOVA, followed by Tukey honestly significant difference test for multiple comparisons. DAPI indicates 4ʹ,6-diamidino-2-phenylindole; α-SMA, α-smooth muscle actin; WGA, wheat germ agglutinin; f/f, flox/flox; Nor, normoxia; SuHx, Sugen5416/hypoxia; and RV, right ventricular.

### E64FC26 Exhibits Therapeutic Effects Against PH in Rats

E64FC26 is a potent PDI pan-inhibitor with an IC_50_ of 16.3 μM against TXNDC5^18^. To assess its safety, histological analyses were performed on major organs from rats treated with E64FC26 for 5 weeks under normoxic conditions. No pathological abnormalities were observed in the kidneys, liver, or spleen, with preserved tissue architecture and no evidence of necrosis or inflammatory infiltration (**Figure S2A**). These findings indicate that E64FC26 exhibits a favorable safety profile without causing overt organ or systemic toxicity.

To examine the therapeutic effect of TXNDC5 inhibition in PH, E64FC26 was administered intraperitoneally in SuHx-induced PH rats (**Figure 4A**). E64FC26 administration significantly attenuated the increases of RVSP (**Figure 4B**) and RV/(LV+S) ratio (**Figure 4C**) compared with vehicle-treated rats. E64FC26 reversed SuHx-induced pulmonary arterial muscularization (**Figure 4D**) and wall thickness (**Figure 4E**). In addition, histological analyses revealed that E64FC26 markedly alleviated RV hypertrophy in SuHx-induced PH rats (**Figure 4F**). Echocardiographic assessment further demonstrated improved RV function following E64FC26 treatment (**Figure S2B**).

**Figure 4.**
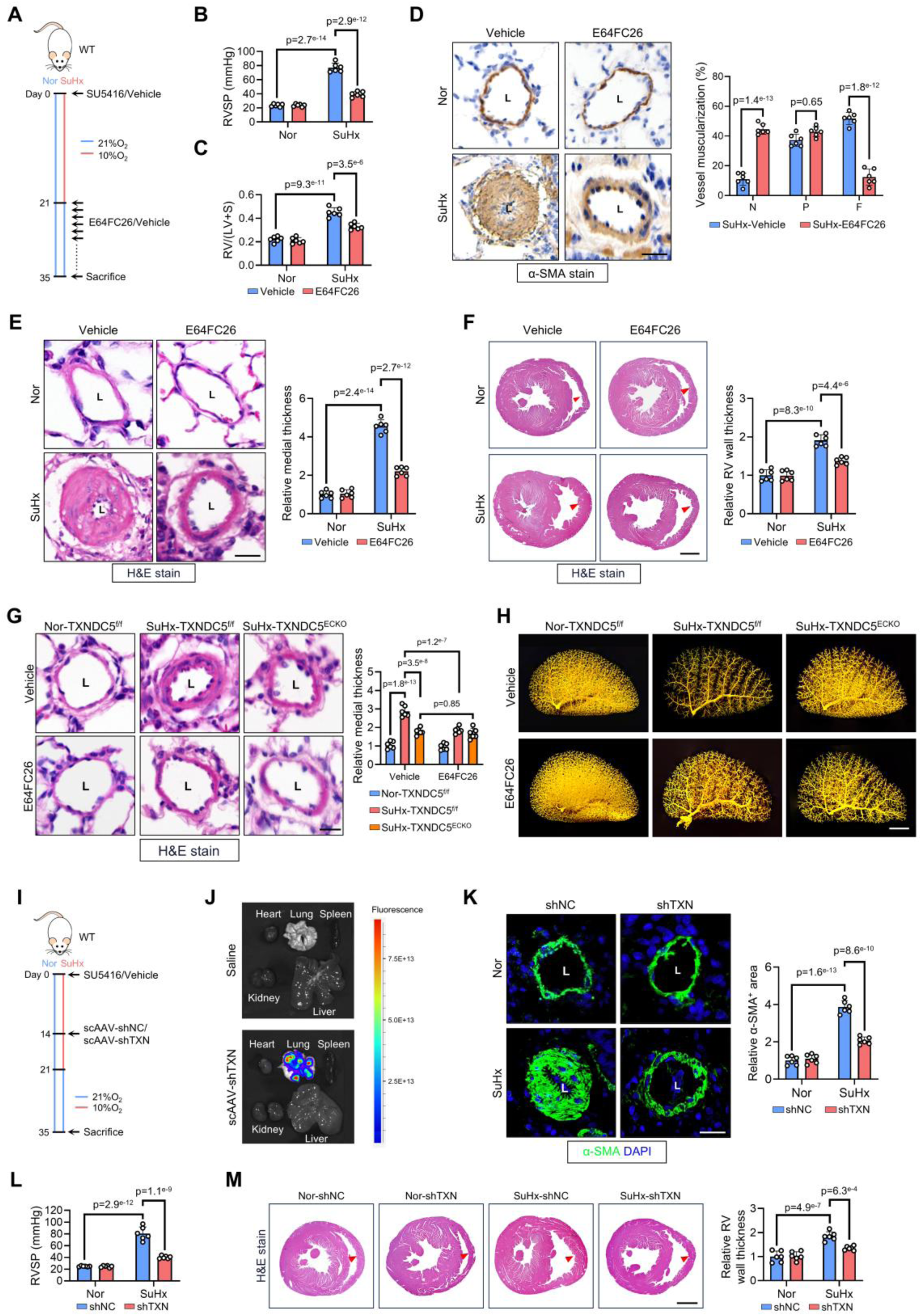
Pharmacological and pulmonary endothelium-targeted inhibition of TXNDC5 ameliorated PH. **A**, Schematic showing the E64FC26 treatment protocol in SuHx-induced PH rats. **B**, Hemodynamic measurements showing right ventricular systolic pressure (RVSP) in vehicle- or E64FC26-treated rats (n=6). **C**, Fulton index [RV/(LV+S)] in rats treated with vehicle or E64FC26 (n=6). **D**, Immunohistochemical staining for α-SMA showing muscularization of distal pulmonary arteries in vehicle- or E64FC26-treated rats (n=6). L indicates lumen. Scale bar=25 μm. **E**, Hematoxylin and eosin (H&E) staining showing distal pulmonary arterial morphology in vehicle- or E64FC26-treated rats (n=6). L indicates lumen. Scale bar=25 μm. **F**, Histological sections of mid-right ventricle showing RV wall thickening in vehicle- or E64FC26-treated rats (n=6). Scale bar=2 mm. **G**, H&E staining showing distal pulmonary arterial morphology in TXNDC5^f/f^ and TXNDC5^ECKO^ mice treated with vehicle or E64FC26 (n=6). L indicates lumen. Scale bar=25 μm. **H**, Lung angiography images showing distal pulmonary perfusion. Scale bar=2 mm. **I**, Schematic showing pulmonary endothelium-targeted gene therapy using self-complementary adeno-associated virus (scAAV) encoding shRNA against TXNDC5 (shTXN) or control shRNA (shNC). **J**, IVIS imaging showing lung fluorescence signals in scAAV-treated rats. **K**, Immunofluorescence staining showing α-SMA (green) positive area in distal pulmonary arteries in shNC- or shTXN-treated rats (n=6). Nuclei were counterstained with DAPI (blue). L indicates lumen. Scale bar=25 μm. **L**, Hemodynamic measurements showing RVSP in shNC- or shTXN-treated rats (n=6). **M**, H&E staining of mid-right ventricle sections showing RV wall morphology in shNC- or shTXN-treated rats (n=6). Scale bar=2 mm. Data represent the mean±SD. Comparisons of parameters were performed with 2-way ANOVA, followed by Tukey honestly significant difference test for multiple comparisons.α-SMA indicates α-smooth muscle actin; N, non-muscularized; P, partially muscularized; F, fully muscularized; shRNA, short hairpin RNA; f/f, flox/flox; Nor, normoxia; SuHx, Sugen5416/hypoxia; and RV, right ventricular.

To determine whether the therapeutic effects of E64FC26 in PH are mediated primarily through TXNDC5 rather than off-target inhibition of other PDI family members, TXNDC5^ECKO^ mice were treated with E64FC26 under SuHx conditions (**Figure S2C**). E64FC26 did not provide additional protection in TXNDC5 endothelial-deficient mice, as distal pulmonary arterial remodeling remained unchanged compared with vehicle-treated TXNDC5^ECKO^ mice (**Figure 4G**). Lung angiography further showed no improvement in distal pulmonary perfusion following E64FC26 treatment (**Figure 4H**). Consistently, E64FC26 failed to attenuate RVSP elevation (**Figure S2D**), RV hypertrophy (**Figure S2E**), and RV dysfunction (**Figure S2F**) in SuHx-treated TXNDC5^ECKO^ mice. These findings indicate that the therapeutic efficacy of E64FC26 in PH is largely dependent on endothelial TXNDC5, supporting pharmacological inhibition of TXNDC5 as a promising strategy for PH treatment.

### Pulmonary Endothelium-Targeted TXNDC5 Gene Therapy Ameliorates PH

We further established a pulmonary endothelium-targeted gene therapy approach to enhance organ specificity. Using a self-complementary adeno-associated virus (scAAV) vector encoding shRNA against TXNDC5 (shTXN), we achieve rapid and endothelial-specific gene silencing *in vivo*. The vector was administered via the tail vein to SuHx-induced PH rats to selectively suppress TXNDC5 in lung ECs (**Figure 4I**). *Ex vivo* imaging confirmed efficient lung targeting (**Figure 4J**), and Western blotting showed marked TXNDC5 suppression in primary rat lung ECs (**Figure S2G**).

Treatment with shTXN significantly attenuated SuHx-induced pulmonary arterial remodeling compared with the control vector (shNC) (**Figure 4K**). Hemodynamic measurements showed reduced RVSP following shTXN administration (**Figure 4L**). Moreover, morphological and histological analyses showed that shTXN decreased the RV/(LV+S) ratio (**Figure S2H**) and alleviated RV wall thickening (**Figure 4M**). Consistently, echocardiographic assessment revealed improved RV function (**Figure S2I**). Collectively, these results demonstrate that endothelial-targeted TXNDC5 silencing effectively mitigates pulmonary vascular remodeling and right heart dysfunction in PH, highlighting the therapeutic potential of TXNDC5 inhibition.

### TXNDC5 Promotes Hypoxia-Induced Dysfunction in PAECs

Exposure to hypoxia significantly increased the TXNDC5 protein and mRNA levels in PAECs (**Figure S3A-B**), predominantly in the cytoplasm (**Figure S3C**). Silencing TXNDC5 with small-interfering RNA oligonucleotides (siTXNDC5) suppressed proliferation (**Figure S3D**) and migration (**Figure S3E**) of PAECs under both normoxic and hypoxic conditions, and prevented G1-to-S phase transition (**Figure S3F**). Conversely, TXNDC5 overexpression via lentivirus (LV-TXNDC5) disrupted cytoskeletal architecture, inducing prominent stress fibers and diffuse paxillin distribution (**Figure S3G-H**). Taken together, TXNDC5 regulates important functional abnormalities associated with PH pathogenesis, with disruption of cytoskeletal architecture in PAECs.

### HIF-2α Transcriptionally Activates TXNDC5 to Drive Pulmonary Hypertension

As an endothelial-specific hypoxia-responsive transcription factor, HIF-2α has been considered a critical regulator of pulmonary vascular remodeling in PH^19^. We examined the HIF-2α activity based on the single-cell RNA sequencing dataset (GSE273062) and found that it was markedly elevated in ECs from SuHx-treated rat lungs compared with those from normoxic controls (**Figure 5A**). Consistently, immunofluorescence staining of remodeled pulmonary arteries from patients with PH revealed pronounced nuclear accumulation of HIF-2α within ECs, accompanied by robust expression of TXNDC5 (**Figure 5B**). These findings implicate HIF-2α as a transcriptional activator of TXNDC5 in pulmonary arterial ECs during PH. To further determine whether TXNDC5 is a direct transcriptional target of HIF-2α, we analyzed the promoter region of TXNDC5 and identified two putative HIF-2α binding sites (BS1 and BS2) (**Figure 5C**). Chromatin immunoprecipitation (ChIP) assays confirmed that HIF-2α bound to both sites in the TXNDC5 promoter (**Figure 5D**). Consistently, dual-luciferase reporter assays demonstrated that mutation of either binding site significantly reduced TXNDC5 promoter activity, whereas simultaneous mutation of both sites caused a further decrease (**Figure 5E**). These results indicate that BS1 and BS2 cooperatively mediate HIF-2α-driven transcriptional activation of TXNDC5.

**Figure 5.**
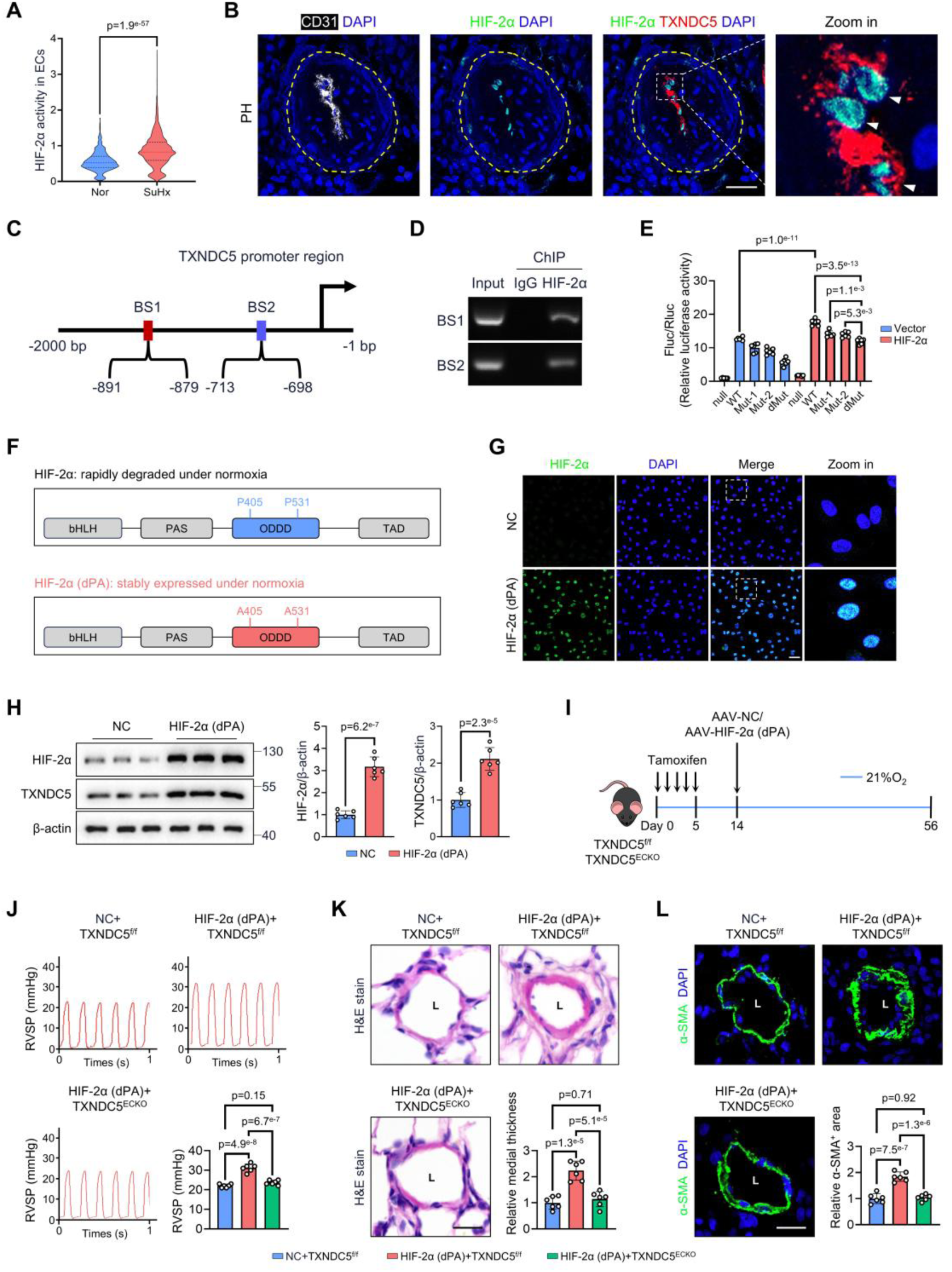
HIF-2α transcriptionally activated TXNDC5 to drive PH. **A**, Single-cell RNA sequencing analysis showing HIF-2α activity inferred from the expression of its target genes in endothelial cells (ECs) from normoxic (Nor) and SuHx-treated rat lungs. **B**, Immunofluorescence staining of pulmonary arteries from patients with PH showing CD31 (white), HIF-2α (green), and TXNDC5 (red). Nuclei were counterstained with DAPI (blue). Scale bar=25 μm. **C**, Schematic illustration showing two putative HIF-2α binding sites (BS1 and BS2) in the TXNDC5 promoter region. **D**, Chromatin immunoprecipitation (ChIP) followed by polymerase chain reaction analysis showing HIF-2α binding to the BS1 and BS2 regions of the TXNDC5 promoter. **E**, Dual-luciferase reporter assay showing TXNDC5 promoter activity with mutation of the HIF-2α binding sites [Mut-1, Mut-2, or double mutant (dMut)] (n=6). **F**, Schematic of HIF-2α (dPA) mutant harboring double Pro405Ala/Pro531Ala substitutions that enable stable expression under normoxia. **G**, Immunofluorescence staining showing HIF-2α (green) in PAECs after lentiviral delivery of HIF-2α (dPA). Nuclei were counterstained with DAPI (blue). Scale bar=50 μm. **H**, Western blot analysis showing HIF-2α and TXNDC5 expression in PAECs after HIF-2α overexpression (n=6). **I**, Schematic showing the experimental design for endothelial-specific overexpression of HIF-2α (dPA) in TXNDC5^f/f^ and TXNDC5^ECKO^ mice. **J**, Hemodynamic measurements showing right ventricular systolic pressure (RVSP) in TXNDC5^f/f^ and TXNDC5^ECKO^ mice after endothelial HIF-2α overexpression (n=6). **K**, Hematoxylin and eosin (H&E) staining showing distal pulmonary arterial morphology in TXNDC5^f/f^ and TXNDC5^ECKO^ mice (n=6). L indicates lumen. Scale bar=25 μm. **L**, Immunofluorescence staining for α-SMA (green) showing the α-SMA-positive area in pulmonary arteries. Nuclei were counterstained with DAPI (blue). L indicates lumen. Scale bar=25 μm. Data represent the mean±SD. Comparisons of parameters were performed with the 2-tailed Mann-Whitney *U* test (**A**) or unpaired 2-tailed Student *t*-test (**H**) or 1-way (**J-L**) or 2-way (**E**) ANOVA, followed by the Tukey honestly significant difference test for multiple comparisons. Nor indicates normoxia; NC, negative control; α-SMA, α-smooth muscle actin; bHLH, basic helix-loop-helix; PAS, Per-Arnt-Sim motifs; ODDD, oxygen-dependent degradation domain; and TAD, transcriptional activation domain.

Next, to mimic sustained HIF-2α activation independent of hypoxia, we utilized a HIF-2α (dPA) mutant in which Pro405 and Pro531 were substituted with alanine (A)^20^, allowing it to escape Von Hippel-Lindau-mediated degradation and stable expression under normoxic conditions (**Figure 5F**). Immunofluorescence staining showed that lentiviral delivery of HIF-2α (dPA) resulted in prominent nuclear accumulation of HIF-2α in PAECs (**Figure 5G**). Quantitative polymerase chain reaction (qPCR) analysis further demonstrated that HIF-2α overexpression significantly increased the mRNA levels of TXNDC5, together with induction of canonical HIF target genes, including vascular endothelial growth factor A (VEGFA) and endothelin-1 (EDN1). (**Figure S4A**). Western blot analysis confirmed robust HIF-2α overexpression and a concomitant increase in TXNDC5 protein levels (**Figure 5H**). Moreover, HIF-2α silencing abolished the hypoxia-induced upregulation of TXNDC5, indicating that HIF-2α was required for hypoxia-driven TXNDC5 induction (**Figure S4B**).

Next, to assess the *in vivo* functional importance of the HIF-2α-TXNDC5 axis, we overexpressed endothelial HIF-2α (dPA) in both TXNDC5^f/f^ and TXNDC5^ECKO^ mice (**Figure 5I**). Notably, without hypoxia exposure, overexpression of HIF-2α (dPA) was sufficient to induce a mild PH phenotype in TXNDC5^f/f^ mice, as evidenced by elevated right RVSP, increased pulmonary arterial remodeling, and RV hypertrophy. Importantly, these pathological changes were largely abolished in TXNDC5^ECKO^ mice, demonstrating that endothelial TXNDC5 is required for HIF-2α-driven pulmonary vascular remodeling and PH development (**Figure 5J-L** and **Figure S4C**). These findings indicate that TXNDC5 is a critical downstream mediator of HIF-2α-driven pathological remodeling in the pulmonary vasculature.

### TXNDC5 Regulates ECM Expression and Homeostasis

To explore the downstream molecular mechanisms of TXNDC5, we reclustered ECs from the previously single-cell RNA-seq dataset, identifying three transcriptionally distinct subpopulations (**Figure 6A-B**). Cluster 1 was annotated as homeostatic gas-exchange homeostatic ECs, characterized by genes associated with capillary specialization and endothelial quiescence, including *Ca4*, *Cldn5*, *Apln*, *Kdr*, and *Selenop*. The second subset exhibited a transcriptional profile indicative of cytoskeletal remodeling and ECM interaction, with prominent expression of *Myo10*, *Itga1*, *Col4a1*, *Fabp4*, and *Adamts9*, and was defined as remodeling-primed transitional ECs (**Figure 6C**). Cluster 3 displayed robust upregulation of ECM components, including *Fn1*, *Vcan*, *Bgn*, and *Ccn2*, and functional enrichment analysis of its top 200 marker genes further confirmed a significant association with ECM-related terms, such as extracellular region and ECM (**Figure 6D**). Notably, TXNDC5 expression was selectively enriched in this subset, defining a TXNDC5^high^ ECM-producing ECs population (**Figure 6E-F**). Trajectory inference analysis revealed a progressive transition from the gas-exchange homeostatic ECs toward the TXNDC5^high^ ECM-producing ECs phenotype (**Figure 6G**). Collectively, these data suggest that PH is associated with the emergence of a distinct endothelial subpopulation characterized by high TXNDC5 expression and linked to ECM remodeling.

**Figure 6.**
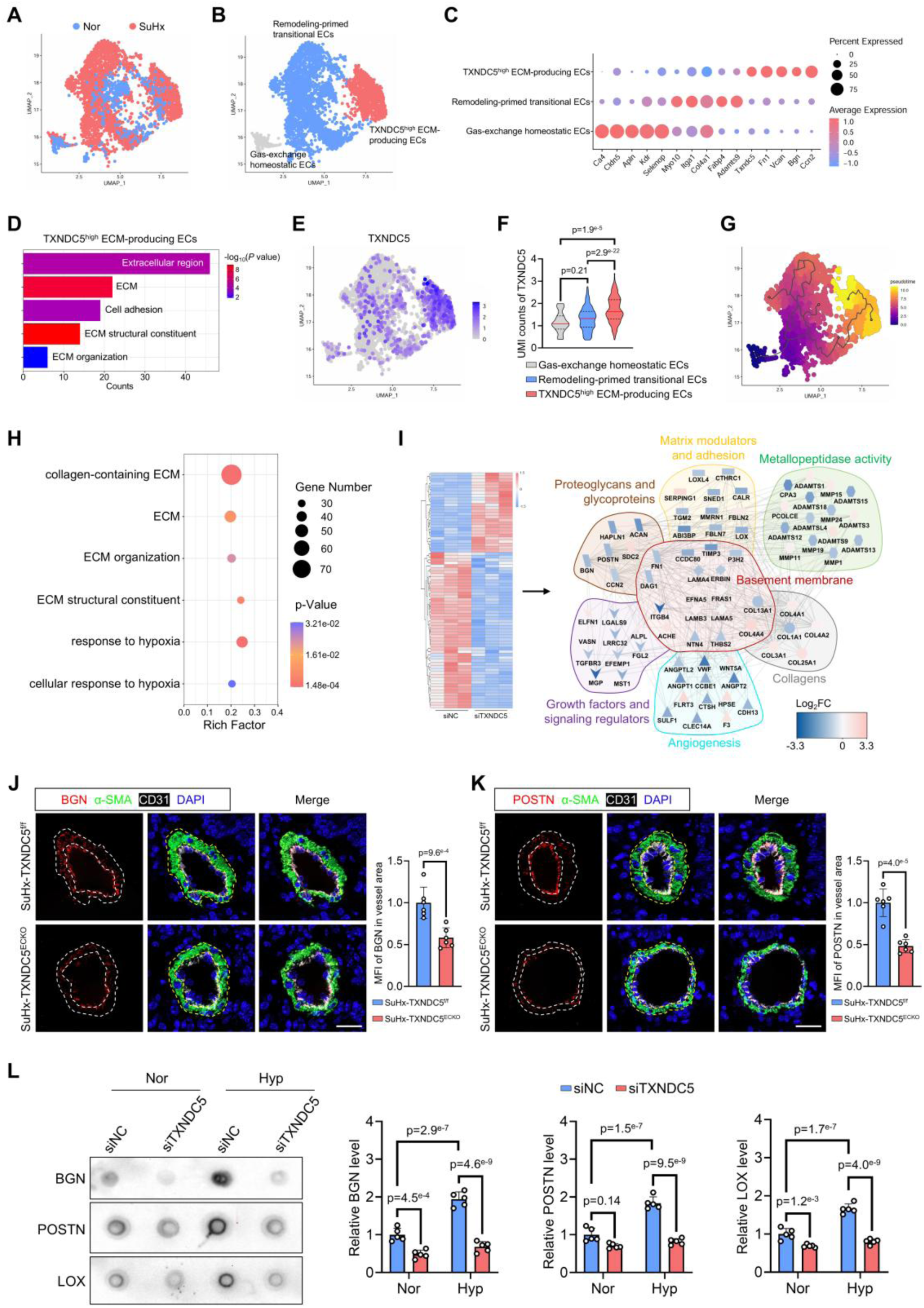
TXNDC5 regulated ECM protein expression and deposition. **A** and **B**, Reclustering analysis of endothelial cells (ECs) from the single-cell RNA-seq dataset, showing three transcriptionally distinct subpopulations in UMAP plots with cluster distribution. **C**, Dot plot showing representative markers for each endothelial population. **D**, Gene ontology enrichment analysis showing the top 200 marker genes in TXNDC5^high^ ECM-producing ECs. **E**, UMAP feature plot showing the distribution of TXNDC5 expression across distinct endothelial clusters. **F,** Quantitative analysis of unique molecular identifier (UMI) counts showing TXNDC5 expression across the three endothelial subpopulations. **G**, Pseudotime trajectory analysis showing the inferred transitional relationship among cell states. **H**, Bubble chart showing the Gene ontology analysis of bulk RNA sequencing data from PAECs transfected with siTXNDC5 or siNC under hypoxia exposure (n=3). **I**, Heatmap showing differentially expressed ECM proteins, together with a PPI network showing functional subgroup classification of the ECM proteins. **J** and **K**, Immunofluorescence staining showing biglycan (BGN) and periostin (POSTN) deposition in pulmonary arteries from TXNDC5^f/f^ and TXNDC5^ECKO^ mice after SuHx exposure (n=6). Nuclei were counterstained with DAPI (blue). Scale bar=25 μm. MFI indicates mean fluorescence intensity. **L**, Dot blot analysis showing BGN, POSTN, and lysyl oxidase (LOX) in supernatants from PAECs (n=5). Data represent the mean±SD. Comparisons of parameters were performed with the unpaired 2-tailed Student *t*-test (**J** and **K**) or 1-way (**F**) or 2-way ANOVA (**L**), followed by Tukey honestly significant difference test for multiple comparisons. ECM indicates extracellular matrix; PPI, protein-protein interaction; Nor, normoxia; Hyp, hypoxia; SuHx, Sugen5416/hypoxia; si, small interfering; and α-SMA, α-smooth muscle actin.

Next, we performed a bulk RNA sequencing (RNA-seq) analysis on PAECs transfected with siRNA targeting TXNDC5 under hypoxia exposure (**Supplemental Material 3**). Gene ontology analysis indicated that the silencing of TXNDC5 resulted in altered expression of genes mainly enriched in “collagen-containing ECM”, “ECM”, “ECM organization”, “ECM structural constituent”, “response to hypoxia”, and “cellular response to hypoxia” (**Figure 6H**). We selected the differentially expressed ECM genes and then constructed a protein-protein interaction (PPI) network (**Figure 6I** and **Supplemental Material 4**). The network revealed that TXNDC5-affected ECM proteins were clustered into different functional subgroups, including “proteoglycans and glycoproteins”, “collagens”, “matrix modulators and adhesion”, “metallopeptidase activity”, “basement membrane”, “growth factors and signaling regulators”, and “angiogenesis”. Immunofluorescence staining demonstrated markedly increased periostin (POSTN) and biglycan (BGN) deposition in pulmonary arteries of TXNDC5^f/f^ mice compared with TXNDC5^ECKO^ mice after SuHx exposure (**Figure 6J-K**). Western blot analysis also confirmed that TXNDC5 silencing reversed the abnormal expression of ECM proteins, including collagen type I alpha 1 chain (COL1A1), transglutaminase 2 (TGM2), fibronectin 1 (FN1), POSTN, and lysyl oxidase (LOX) induced by hypoxia treatment in PAECs (**Figure S4D**). These data indicate that endothelial TXNDC5 deficiency attenuates ECM accumulation. To further investigate whether TXNDC5 regulates ECM secretion, we collected the supernatants from PAECs. Dot blot analysis revealed that the silence of TXNDC5 markedly reduced the secretion of BGN, POSTN, and LOX (**Figure 6L**). Collectively, these data highlight the key role TXNDC5 played in regulating ECM homeostasis.

HIF signaling critically drives ECM remodeling^19^. We found that HIF-2α activity was markedly increased in TXNDC5^high^ ECM-producing ECs (**Figure S4E**). Overexpression of HIF-2α (dPA) in PAECs induced upregulation of representative ECM genes (**Figure S4F**). This increase was further confirmed at the protein level (**Figure S4G**). To further determine whether endothelial TXNDC5 is required for HIF-2α-induced ECM remodeling, we performed immunofluorescence staining of pulmonary arteries. Overexpression of HIF-2α resulted in pronounced deposition of BGN and POSTN in the pulmonary arteries of TXNDC5^f/f^ mice, whereas markedly attenuated in TXNDC5^ECKO^ mice (**Figure S4H-I**). These results indicate that endothelial TXNDC5 is required for HIF-2α-driven ECM accumulation during pulmonary vascular remodeling.

### BGN Is a Promising Biomarker and Therapeutic Target for Pulmonary Hypertension

Given the complexity of the ECM network, we aimed to identify key ECM nodes that are directly regulated by TXNDC5 and critically influence endothelial cell function. We performed a computational analysis of the PPI network by integrating an ECM protein dataset with an endothelial cell function-related protein dataset. This approach allowed us to construct a comprehensive PPI network and further refine it into a core subnetwork using degree-based filtering (**Figure 7A** and **Supplemental Material 5**). Focusing on ECM nodes from the core PPI network and ECM genes enriched in TXNDC5^high^ ECM-producing ECs, we pinpointed BGN as a central hub (**Figure 7B**). This finding positions BGN as a key mediator, capable of orchestrating the broader ECM network to modulate endothelial cell function.

**Figure 7.**
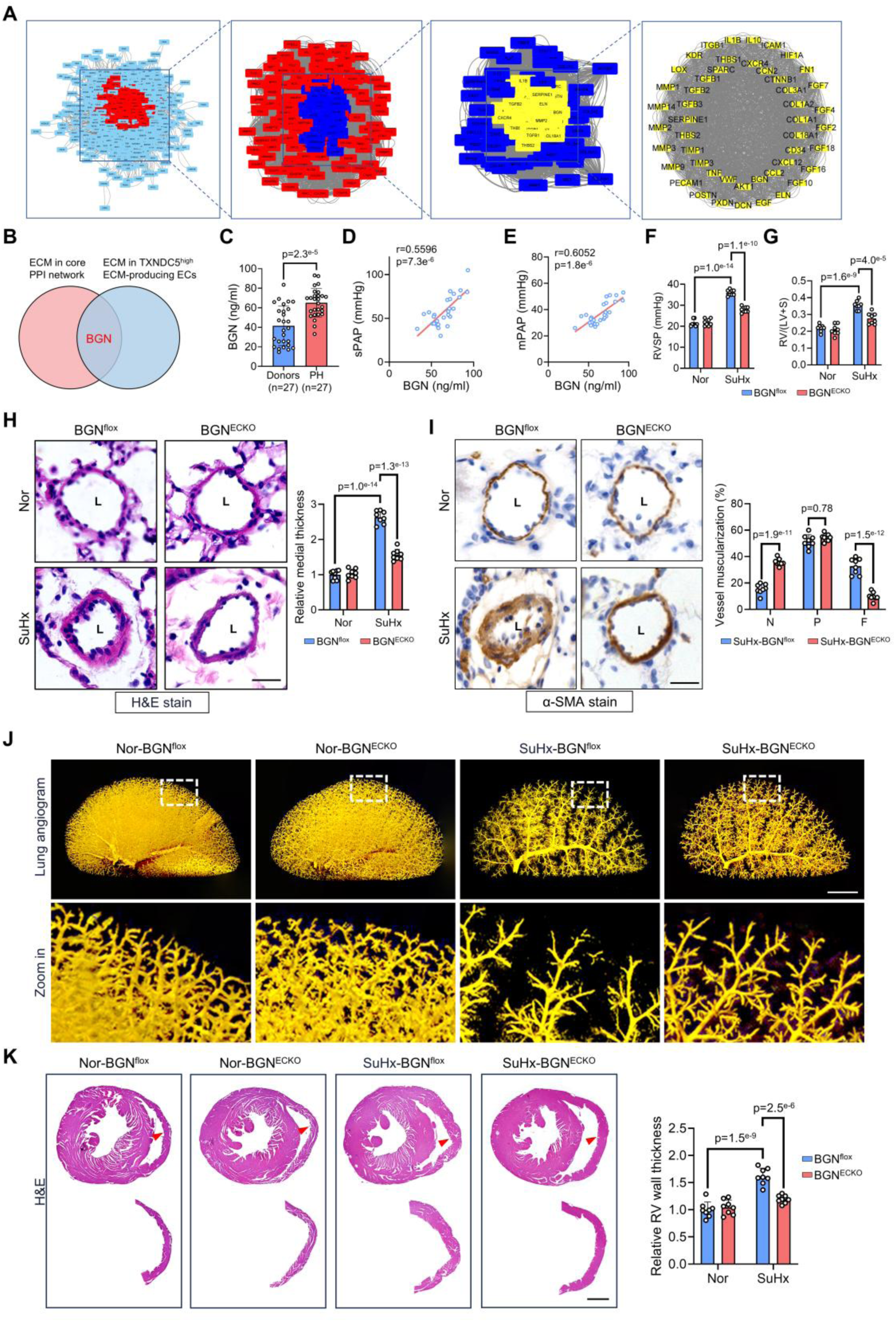
BGN served as a biomarker and therapeutic target of PH. **A**, Progressive PPI network analysis showing interaction between ECM genes and endothelial function-related genes, with each network building on the core genes of the previous one. **B**, Venn diagram illustrating the overlap between ECM proteins in the final core PPI network and ECM proteins enriched in TXNDC5^high^ ECM-producing ECs. **C,** Enzyme-linked immunosorbent assay showing the serum BGN concentrations in PH patients and non-PH donors (n=27 per group). **D-E**, Linear regression analysis showing the correalation between serum BGN levels and sPAP or mPAP in PH patients (n=27). **F**, Hemodynamic measurements showing right ventricular systolic pressure (RVSP) in BGN^flox^ and BGN^ECKO^ mice under the indicated conditions (n=8). **G**, Fulton index [RV/(LV+S)] in BGN^flox^ and BGN^ECKO^ mice (n=8). **H**, Hematoxylin and eosin (H&E) staining showing distal pulmonary arterial morphology in BGN^flox^ and BGN^ECKO^ mice. L indicates lumen. Scale bar=25 μm. **I**, Immunohistochemical staining for α-SMA showing muscularization of pulmonary arteries (n=8). L indicates lumen. Scale bar=25 μm. **J**, Lung angiogram displaying distal pulmonary vascular filling in BGN^flox^ and BGN^ECKO^ mice. Scale bar=2 mm. **K**, H&E staining of mid-right ventricle sections showing RV wall thickening in BGN^flox^ and BGN^ECKO^ mice (n=8). Scale bar= 1 mm. Data represent the mean±SD. Comparisons of parameters were performed with the 2-tailed Mann-Whitney *U* test (**C**), Pearson correlation test (**D** and **E**), or 2-way ANOVA (**F-I**, and **K**), followed by Tukey honestly significant difference test for multiple comparisons. α-SMA indicates α-smooth muscle actin; DAPI, 4ʹ,6-diamidino-2-phenylindole; Nor, normoxia; SuHx, Sugen5416/hypoxia; N, non-muscularized; P, partially muscularized; F, fully muscularized; RV, right ventricular; sPAP, systolic pulmonary arterial pressure; and mPAP, mean pulmonary arterial pressure.

We next investigated the role of BGN in PH. BGN protein levels were increased in lung biopsies from patients with PH (**Figure S5A**). Similarly, BGN was upregulated in the lungs of SuHx-treated mice (**Figure S5B**). As a circulating protein, we also investigated whether serum BGN levels can reflect the severity of PH. Surprisingly, a markedly increased serum BGN level was found in patients with PH compared with the donor controls (**Figure 7C**). Serum BGN levels exhibited a significant correlation with systolic pulmonary artery pressure (sPAP) and mean pulmonary artery pressure (mPAP) (**Figure 7D-E**), indicating its potential as a biomarker. To evaluate the role of BGN in PH generation, we generated endothelial-specific BGN knockout mice (BGN^ECKO^) (**Figure S5C**). Notably, SuHx-induced increases in RVSP and RV/(LV+S) were significantly attenuated in BGN^ECKO^ mice compared with BGN^flox^ controls (**Figure 7F-G**). Consistently, BGN^ECKO^ mice showed reduced pulmonary arterial medial wall thickness and muscularization (**Figure 7H-I**). Lung angiogram also revealed the better-preserved lung vasculature in BGN^ECKO^ mice (**Figure 7J**). Histological assessment of the RV showed a significant reduction in hypertrophy in BGN^ECKO^ mice treated with SuHx (**Figure 7K**). Echocardiography further demonstrated that SuHx-induced decreases in PAT/PET, TAPSE, RVFAC, and increases in RVFWT were less pronounced in BGN^ECKO^ mice than in BGN^flox^ mice (**Figure S5D**). Together, our data suggest that endothelial BGN plays an important role in PH development.

### TXNDC5 Assists BGN Protein Folding to Drive Pulmonary Hypertension

To elucidate how TXNDC5 regulates BGN, we first performed Western blot analysis to validate the regulatory relationship. BGN expression was markedly reduced in primary lung ECs from TXNDC5^ECKO^ mice (**Figure S5E**), whereas it was elevated in lung ECs from AAV-TXN-treated mice (**Figure S5F**), confirming that TXNDC5 positively regulates BGN. To assess the interaction between TXNDC5 and BGN, we first examined their co-localization in single-cell data, which revealed enrichment of BGN in TXNDC5^high^ ECM-producing ECs (**Figure S5G**). Subsequent immunostaining confirmed strong co-localization of TXNDC5 and BGN in PAECs (**Figure 8A**). Co-immunoprecipitation (co-IP) analysis also showed that TXNDC5 bound to BGN, and the binding was enhanced upon hypoxia exposure (**Figure 8B**). To identify the protein domain of TXNDC5 that was involved in the interaction with BGN, we performed molecular docking analysis and revealed that multiple sites of TXNDC5 (Cys89, Ser125, Gly351, and His352) might be involved in the interaction with BGN (**Figure 8C**). Thus, we constructed four TXNDC5 mutants (C89A, S125A, G351A, and H352A) and demonstrated that C89A and G351A partially abolished TXNDC5 binding with BGN (**Figure 8D**), suggesting that these amino acid residues might be essential for the binding of TXNDC5 and BGN.

**Figure 8.**
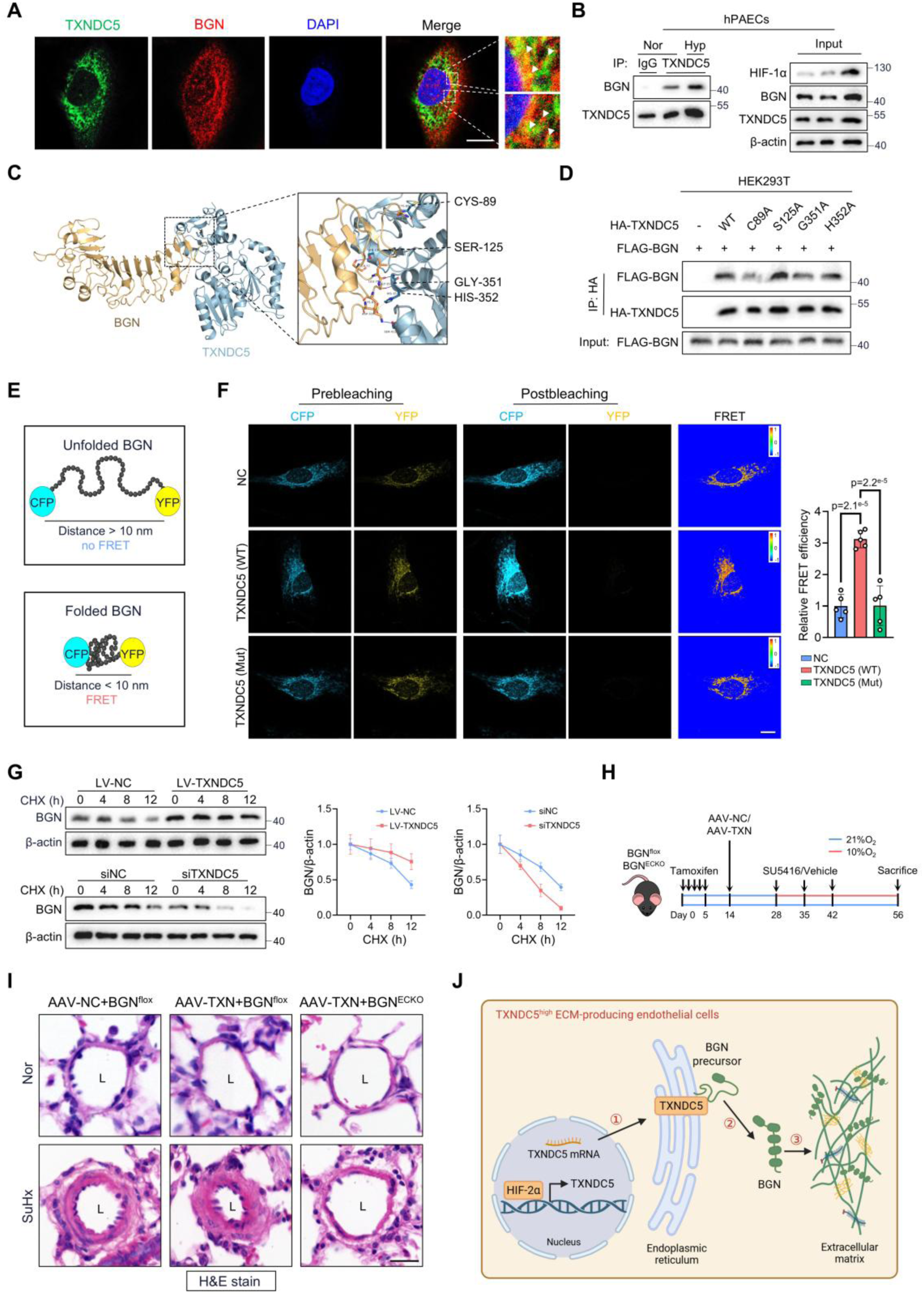
TXNDC5 assisted BGN protein folding to promote PH. **A**, Immunofluorescence staining showing TXNDC5 (green) and BGN (red) localization in PAECs. Nuclei were stained with DAPI (blue). Scale bar=10 μm. **B**, Coimmunoprecipitation (co-IP) analysis showing the interaction between TXNDC5 and BGN in PAECs under both normoxic and hypoxic situations. IgG was used as the negative control. **C**, Molecular docking showing the interaction sites between TXNDC5 and BGN, highlighting Cys89, Ser125, Gly351, and His352 residues of TXNDC5. **D**, Immunoprecipitation with anti-HA beads in HEK293T cells transfected with FLAG-BGN and HA-TXNDC5 mutant plasmids. **E**, Schematic illustration showing the FRET-based assay used to assess BGN protein folding in PAECs. When the BGN protein folds correctly (bottom), the distance between CFP and YFP will be less than 10 nm, resulting in FRET occurring. **F**, FRET-based protein folding assays demonstrating BGN folding efficiency in PAECs expressing wild-type TXNDC5 (TXNDC5 WT) or PDI-inactive TXNDC5 mutant (TXNDC5 Mut) (n=5). Scale bar=10 μm. **G**, Cycloheximide (CHX) chase assays showing BGN protein stability in PAECs with TXNDC5 overexpression or silencing (n=6). **H**, Schematic showing the experimental design for endothelial-specific TXNDC5 overexpression in BGN^flox^ and BGN^ECKO^ mice under SuHx exposure. **I**, Hematoxylin and eosin (H&E) staining showing distal pulmonary arterial morphology in BGN^flox^ and BGN^ECKO^ mice under indicated conditions (n=6). L indicates lumen. Scale bar=25 μm. **J**, Schematic illustration of the HIF-2α-TXNDC5-BGN axis regulating ECM remodeling in TXNDC5^high^ ECM-producing endothelial cells during PH development. Data represent the mean±SD. Comparisons of parameters were performed with the 1-way (**F**) or 2-way (**G**) ANOVA, followed by the Tukey honestly significant difference test for multiple comparisons. Nor indicates normoxia; Hyp, hypoxia; IgG, immunoglobulin G; FRET, fluorescence resonance energy transfer; CFP, cyan fluorescent protein; and YFP, yellow fluorescent protein.

As an endoplasmic reticulum resident protein with three thioredoxin domains containing CGHC motifs, TXNDC5 catalyzes disulfide bond formation^11^. Thus, we hypothesized that TXNDC5 facilitates the folding of BGN, enabling its maturation, signaling, and secretion. To test this, we generated a PDI-inactive TXNDC5 mutant (TXNDC5 Mut) with cysteine-to-alanine substitutions in all three domains. Fluorescence resonance energy transfer (FRET)-based protein folding assays showed that wild-type TXNDC5 (TXNDC5 WT) increased BGN folding efficiency, whereas TXNDC5 Mut did not (**Figure 8E-F**). Moreover, cycloheximide (CHX) chase assays further demonstrated that overexpression of TXNDC5 markedly prolonged the half-life of BGN, whereas TXNDC5 silencing accelerated BGN degradation (**Figure 8G**). These results indicate that TXNDC5 relies on its PDI activity to promote BGN folding and stability.

To assess whether BGN mediates the endothelial TXNDC5 *in vivo*, we overexpressed TXNDC5 in the lung ECs of BGN^ECKO^ and BGN^flox^ mice (**Figure 8H**). AAV-TXN treatment exacerbated pulmonary arterial remodeling in BGN^flox^ mice under SuHx conditions, whereas this effect was largely abolished in BGN^ECKO^ mice (**Figure 8I** and **Figure S5H**). Consistently, AAV-TXN markedly increased RVSP (**Figure S5I**) and aggravated RV hypertrophy in BGN^flox^ mice, whereas both changes were significantly attenuated in BGN^ECKO^ mice (**Figure S5J**). Echocardiographic assessment revealed that AAV-TXN led to pronounced deterioration of RV performance, while these adverse functional changes were largely blunted in BGN^ECKO^ mice (**Figure S5K**). Together, these results indicate that endothelial BGN is required for TXNDC5-mediated PH development, confirming BGN as a key downstream effector of TXNDC5.

## DISCUSSION

PDI family catalyzes disulfide bond formation and rearrangement in the endoplasmic reticulum. Using 4D label-free proteomics, we identified TXNDC5 as a potential contributor to PH. TXNDC5, dominantly expressed in ECs, was also known as endothelial PDI (Endo-PDI)^21^. We found that the silencing of TXNDC5 inhibited the proliferation and migration of PAECs, whereas its overexpression led to changes in the cytoskeletal architecture of PAECs, implying a critical role of TXNDC5 in endothelial dysfunction in PH. *In vivo*, global or endothelial-specific TXNDC5 deletion attenuated, while endothelial TXNDC5 overexpression exacerbated SuHx-induced PH. Pharmacological inhibition of TXNDC5 with E64FC26 alleviated pulmonary arterial remodeling and right ventricular hypertrophy, thereby preventing PH progression in rats. These findings identify TXNDC5 as a promising therapeutic target for PH.

We next examined the molecular mechanisms of the critical role of TXNDC5 in PH development. Accumulating evidence indicates that ECM remodeling, which precedes pulmonary vascular remodeling, is a key driver of PH progression by altering vascular stiffness and resistance^22^. In this study, through single-cell transcriptomic analysis, we identified a distinct endothelial subpopulation characterized by high TXNDC5 expression and a transcriptional signature enriched for ECM production and remodeling (TXNDC5^high^ ECM-producing ECs). Subsequent bulk RNA sequencing and PPI analysis revealed that TXNDC5 broadly modulated ECM components, including proteoglycans and glycoproteins that govern matrix stiffness and elasticity, collagens that provide structural support, matrix modulators and adhesion proteins involved in ECM crosslinking, and metallopeptidases responsible for ECM turnover. TXNDC5 also influenced basement membrane proteins and angiogenesis-related factors, thereby affecting endothelial barrier function and vascular homeostasis. Notably, excessive deposition of BGN and POSTN was observed in the pulmonary arteries of SuHx-treated TXNDC5^f/f^ mice but was markedly reduced in TXNDC5^ECKO^ mice. As key ECM regulators, POSTN and BGN promote matrix organization and mechanotransduction^23–25^, collectively driving vascular stiffening and remodeling. These findings imply that TXNDC5 inhibition shifts the ECM equilibrium toward a less fibrotic and more compliant phenotype, mitigating PH progression by preserving vascular ECM homeostasis.

BGN has been implicated in inflammation, metabolic, musculoskeletal, and oncogenic diseases^26^. In the present study, we identified BGN as a direct binding partner of TXNDC5 and a central downstream mediator to regulate ECM homeostasis in ECs. Mutations at Cys89 and Gly351 within the PDI active domains of TXNDC5 markedly disrupted TXNDC5-BGN binding, indicating that PDI activity is essential for this interaction. Consistently, WT-TXNDC5, but not the PDI-inactive mutant, significantly increased FRET efficiency in the BGN folding assay, demonstrating that TXNDC5 facilitates BGN folding in a PDI-dependent manner. This process is critical for BGN maturation, stability, and secretion. Further investigation revealed that BGN was upregulated in the lungs of PH patients and rodent models. Endothelial-specific deficiency of BGN significantly alleviated PH progression in mice, highlighting endothelial-derived BGN as a key contributor to disease pathogenesis. Importantly, BGN deficiency largely abolished TXNDC5 overexpression-induced distal pulmonary arterial remodeling and RV hypertrophy, establishing BGN as a critical downstream effector of TXNDC5-driven PH. Notably, serum BGN levels were significantly elevated in PH patients and strongly correlated with disease severity, suggesting that BGN may serve as a potential diagnostic biomarker for PH.

Endothelial HIF-2α plays a pivotal role in the pathogenesis of PH^27^. HIF-2α is predominantly localized in ECs and drives sustained transcriptional programs that promote pulmonary vascular remodeling and endothelial dysfunction^28^. In this study, we found that HIF-2α directly binds the promoter region of TXNDC5, driving its transcription. Consistent with previous studies that endothelial loss of prolyl hydroxylase domain protein 2 leads to HIF-2α stabilization and spontaneous PH^29,30^, we demonstrated that endothelial-specific overexpression of HIF-2α *in vivo* induced mile PH phenotypes. Notably, these pathological effects were largely abolished in TXNDC5^ECKO^ mice, demonstrating that TXNDC5 is a critical mediator of HIF-2α-driven pulmonary vascular remodeling. Further studies demonstrated that HIF-2α activity was elevated in TXNDC5^high^ ECM-producing ECs. Using a stabilized HIF-2α (dPA) mutant to mimic sustained HIF-2α activation, we demonstrated that HIF-2α is sufficient to transcriptionally induce TXNDC5 and its downstream ECM effectors. Immunofluorescence revealed pronounced deposition of BGN and POSTN in pulmonary arteries upon HIF-2α overexpression, which was markedly attenuated in endothelial TXNDC5-deficient mice. These findings indicate that TXNDC5 acts as a critical downstream effector linking HIF signaling to pathological ECM remodeling, thereby driving pulmonary vascular remodeling and PH development (**Figure 8J**).

In summary, we identified for the first time the novel role of TXNDC5 in the pathogenesis of PH. Endothelial TXNDC5 functions as a key regulator of ECM homeostasis by promoting the proper folding of BGN. HIF-2α transcriptionally activates TXNDC5, linking hypoxic signaling to pathological ECM accumulation. This study highlights TXNDC5 as a promising therapeutic target for PH and lays the groundwork for future clinical investigations.

## Acknowledgments

We thank Professor John Shyy (University of California, San Diego, USA) for critical reading of the manuscript and valuable advice.

## Supplemental Figures

**Figure S1.**
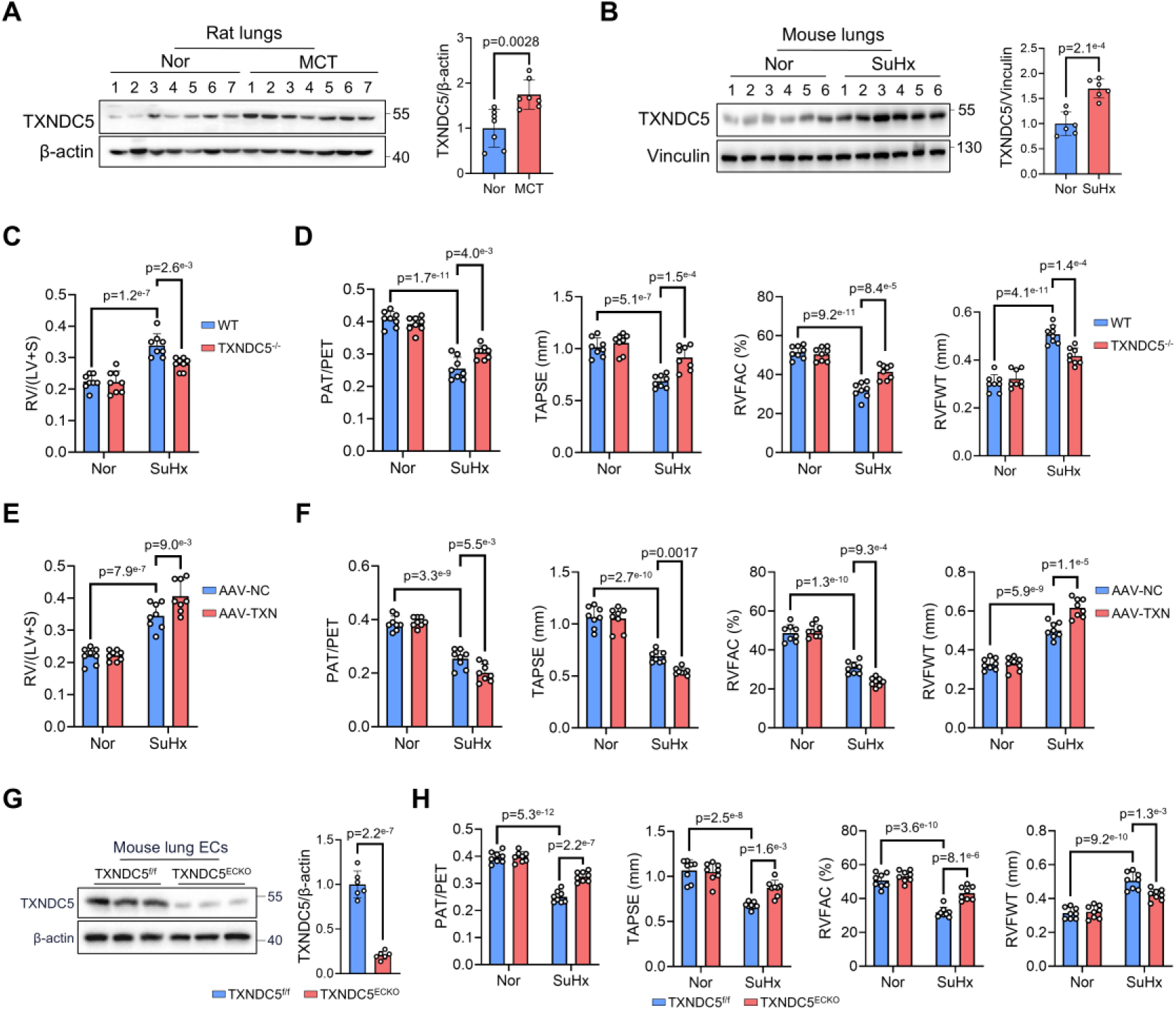
Endothelial TXNDC5 was upregulated and associated with pulmonary vascular remodeling in PH. **A**, Western blot analysis and quantification showing TXNDC5 protein levels in lungs from monocrotaline (MCT)-treated rats and controls (n=7). **B**, Western blot analysis showing TXNDC5 protein levels in lungs from SuHx-exposed mice and normoxic controls (n=6). **C**, Fulton index [RV/(LV+S)] in WT and TXNDC5^-/-^ mice (n=8). **D**, Echocardiography analysis showing right ventricular function parameters in WT and TXNDC5^-/-^ mice (n=8). **E**, RV/(LV+S) in mice receiving pulmonary endothelial TXNDC5 overexpression (AAV-TXN) or control (AAV-NC) (n=8). **F**, Echocardiography analysis showing right ventricular function parameters in AAV-TXN- or AAV-NC-treated mice (n=8). **G**, Western blot analysis showing TXNDC5 expression in lung endothelial cells (ECs) from TXNDC5^ECKO^ and control mice (n=6). **H**, Echocardiography analysis showing right ventricular function parameters in TXNDC5^ECKO^ mice and TXNDC5^f/f^ controls (n=8). Data represent the mean±SD. Comparisons of parameters were performed with the unpaired 2-tailed Student *t*-test (**A-B** and **G**) or 2-way ANOVA (**C**-**F**, and **H**), followed by Tukey honestly significant difference test for multiple comparisons. Nor indicates normoxia; SuHx, Sugen5416/hypoxia; and f/f, flox/flox.

**Figure S2.**
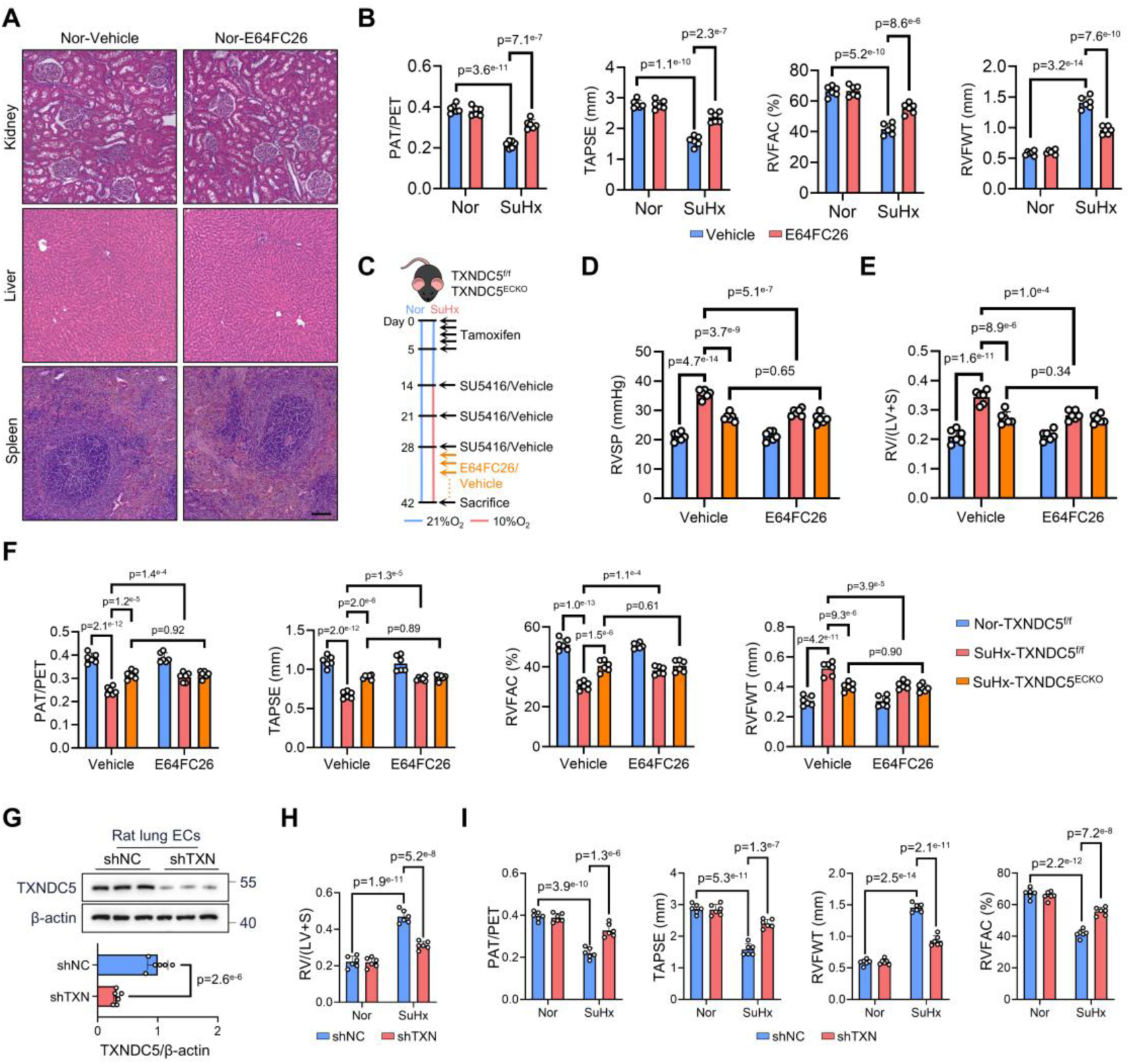
TXNDC5 inhibition showed therapeutic efficacy in PH. **A**, Hematoxylin and eosin (H&E) staining showing kidney, liver, and spleen morphology in normoxic rats treated with E64FC26 or vehicle for 5 weeks. Scale bar=100 μm. **B**, Echocardiography demonstrated improved RV function with E64FC26 treatment (n=6). **C**, Schematic illustrating vehicle or E64FC26 treatment in TXNDC5^f/f^ and TXNDC5^ECKO^ mice. **D-E**, Hemodynamic and morphometric measurements showing right ventricular systolic pressure (RVSP) and the Fulton index [RV/(LV+S)] in TXNDC5^f/f^ and TXNDC5^ECKO^ mice treated with E64FC26 or vehicle (n=6). **F**, Echocardiography analysis showing RV function parameters in TXNDC5^f/f^ and TXNDC5^ECKO^ mice under the indicated conditions (n=6). **G**, Western blot showing TXNDC5 expression in lung endothelial cells (ECs) isolated from rats receiving pulmonary endothelium-targeted self-complementary adeno-associated virus (scAAV) carrying shRNA against TXNDC5 (shTXN) or control shRNA (shNC) (n=6). **H-I**, RV/(LV+S) and echocardiographic RV function parameters in shNC- or shTXN-treated rats (n=6). Data represent the mean±SD. Comparisons of parameters were performed with the unpaired 2-tailed Student *t*-test (**G**) or 2-way ANOVA (**B**, **D-F**, and **H-I**), followed by Tukey honestly significant difference test for multiple comparisons. f/f indicates flox/flox; Nor, normoxia; SuHx, Sugen5416/hypoxia; RV, right ventricular; and shRNA, short hairpin RNA.

**Figure S3.**
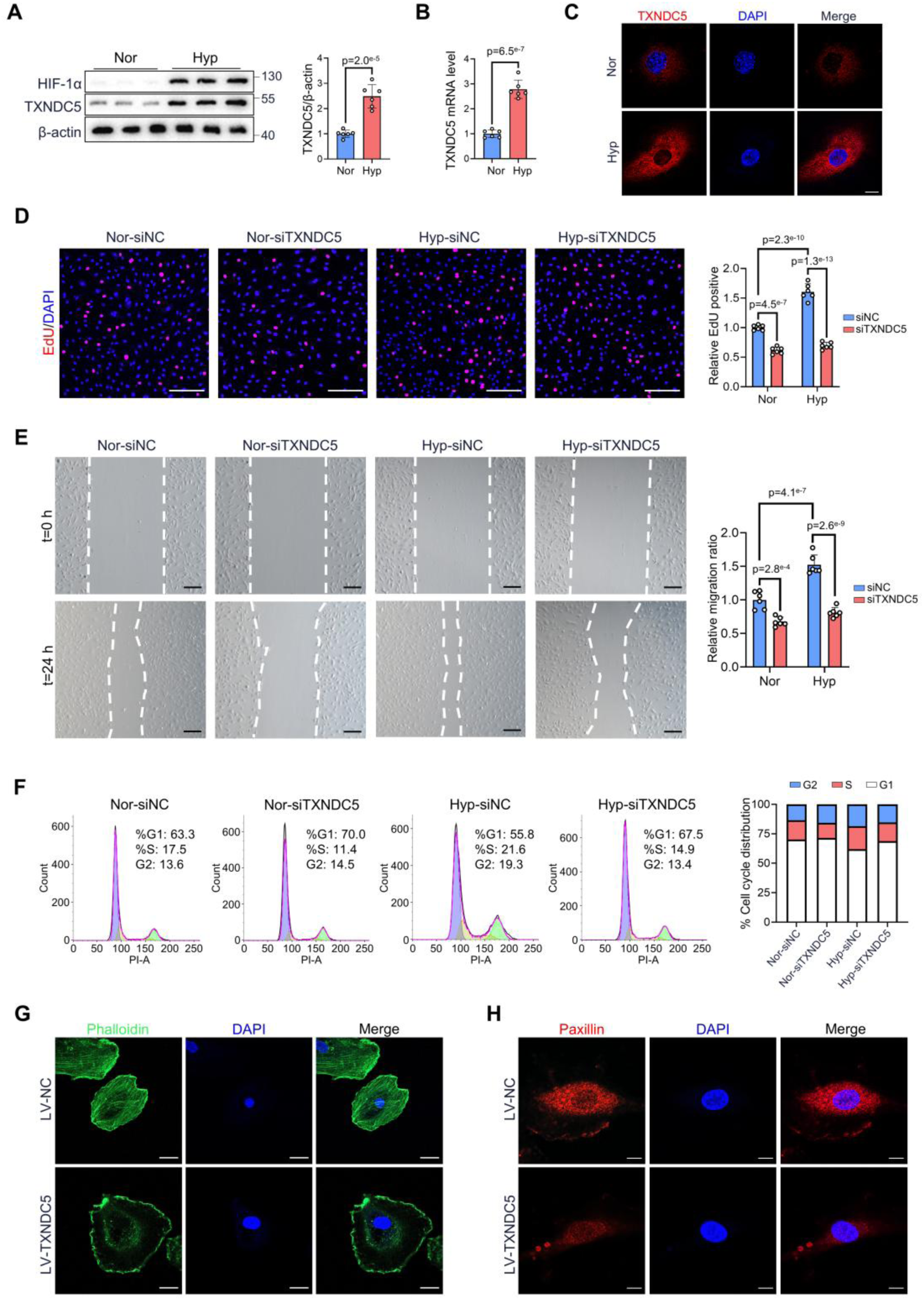
TXNDC5 regulated pulmonary arterial endothelial cell function. **A** and **B**, Western blot and qPCR analyses showing TXNDC5 expression levels in PAECs under normoxia and hypoxia (n=6). **C**, Immunofluorescence staining showing TXNDC5 expression and subcellular localization. Scale bar=10 μm. **D**, Representative EdU staining images and quantification showing PAECs proliferation after TXNDC5 silencing (n=6). Nuclei were counterstained with DAPI (blue). Scale bar=200 μm. **E**, Representative images and group data showing PAECs migration after TXNDC5 silencing under normoxia and hypoxia (n=6). Scale bar=200 μm. **F**, Flow cytometry analysis showing cell cycle distribution in PAECs after TXNDC5 silencing (n=6). **G** and **H**, Immunofluorescence staining showing cytoskeletal organization in PAECs after TXNDC5 overexpression via lentivirus (LV-TXNDC5). Phalloidin (green, scale bar=20 μm) and paxillin (red, scale bar=10 μm) staining were used. Nuclei were counterstained with DAPI (blue). Data represent the mean±SD. Comparisons of parameters were performed with the unpaired 2-tailed Student *t*-test (**A** and **B**) or 2-way ANOVA (**D** and **E**), followed by Tukey honestly significant difference test for multiple comparisons. Nor indicates normoxia; Hyp, hypoxia; si, small interfering; qPCR, quantitative polymerase chain reaction; and EdU, 5-ethynyl-20-deoxyuridine.

**Figure S4.**
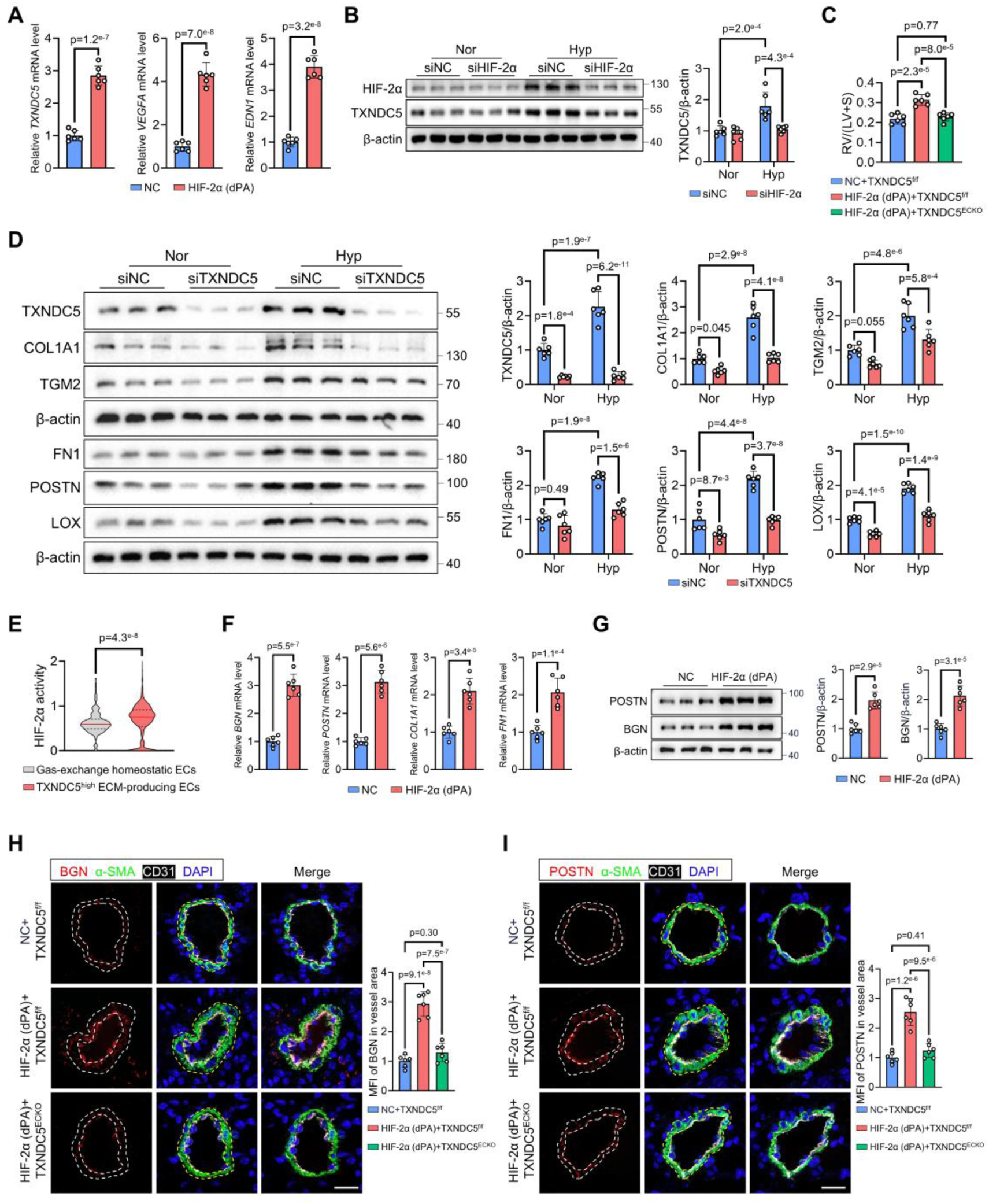
TXNDC5 was required for HIF-2a-driven ECM remodeling. **A**, Quantitative polymerase chain reaction (qPCR) analysis showing mRNA levels of TXNDC5, VEGFA, and EDN1 in PAECs following HIF-2α (dPA) overexpression (n=6). **B**, Western blot analysis showing TXNDC5 protein levels in PAECs under hypoxia with or without HIF-2α silencing (n=6). **C**, Fulton index [RV/(LV+S)] in TXNDC5^f/f^ and TXNDC5^ECKO^ mice after endothelial HIF-2α overexpression (n=6). **D**, Western blot showing ECM protein expression, including COL1A1, TGM2, FN1, POSTN, and LOX, in PAECs following TXNDC5 silencing with or without hypoxia exposure (n=6). **E**, Single-cell RNA-sequencing analysis showing HIF-2α activity in Gas-exchange homeostatic ECs and TXNDC5^high^ ECM-producing ECs. **F**, qPCR showing the mRNA levels of representative ECM genes, including BGN, POSTN, COL1A1, and FN1 in PAECs after HIF-2α overexpression (n=6). **G**, Western blot showing POSTN and BGN protein levels in PAECs (n=6). **H** and **I,** Immunofluorescence staining showing BGN and POSTN in pulmonary arteries of TXNDC5^f/f^ and TXNDC5^ECKO^ mice after endothelial HIF-2α (dPA) overexpression (n=6). Data represent the mean±SD. Comparisons of parameters were performed with the unpaired 2-tailed Student *t*-test (**A, F, and G**) or 2-tailed Mann-Whitney *U* test (**E**) or one-way (**C**, **H,** and **I**) or 2-way ANOVA (**D**), followed by Tukey honestly significant difference test for multiple comparisons. Nor indicates normoxia; Hyp, hypoxia; SuHx, Sugen5416/hypoxia; si, small interfering; and ECs, endothelial cells.

**Figure S5.**
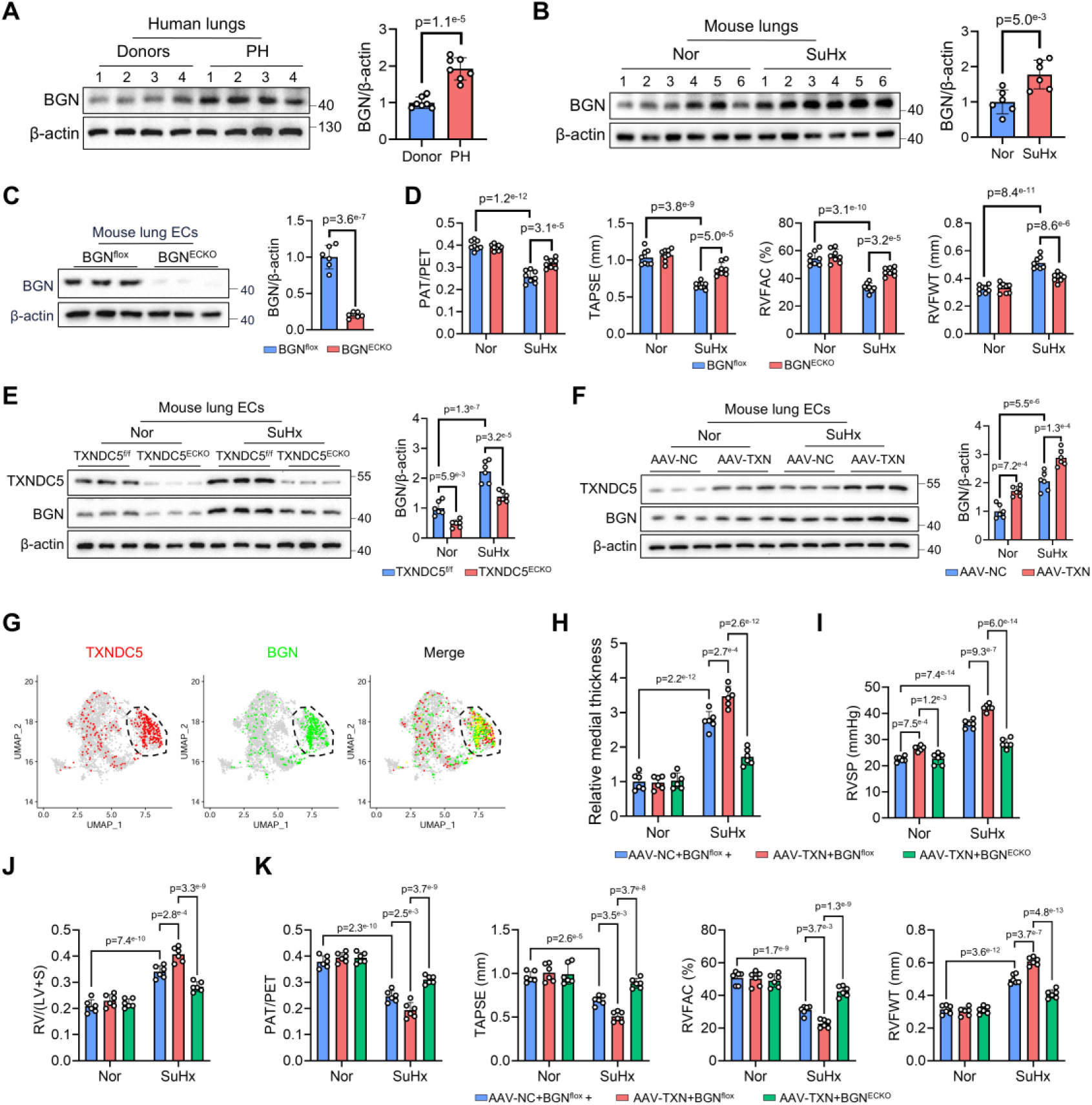
BGN mediated TXNDC5-driven ECM regulation and PH development. **A** and **B**, Western blots showing BGN protein levels in lung tissue from patients with PH (n=7) and from SuHx-treated mice (n=6). **C**, Western blot analysis showing BGN protein levels in isolated lung endothelial cells (ECs) from BGN^ECKO^ mice and BGN^flox^ controls (n=6). **D**, Echocardiographic assessment indicated that SuHx-induced right ventricular dysfunction was attenuated in BGN^ECKO^ mice compared with BGN^flox^ controls (n=6). **E**, Western blot analysis showing BGN protein levels in lung ECs from TXNDC5^f/f^ and TXNDC5^ECKO^ mice under normoxia or SuHx conditions (n=6). **F,** Western blot analysis showing BGN expression in lung ECs following endothelial TXNDC5 overexpression (AAV-TXN) (n=6). **G**, UMAP feature plot showing the distribution of TXNDC5 and BGN expression in TXNDC5^high^ ECM-producing ECs. **H**, Quantification of pulmonary arterial medial wall thickness based on hematoxylin and eosin (H&E) staining in BGN^flox^ and BGN^ECKO^ mice following endothelial TXNDC5 overexpression (n=6). **I**, Measurement of right ventricular systolic pressure (RVSP) in BGN^flox^ and BGN^ECKO^ mice following endothelial TXNDC5 overexpression (n=6). **J**, Fulton index [RV/(LV+S)] in BGN^flox^ and BGN^ECKO^ mice (n=6). **K**, Echocardiographic analysis showing right ventricular function parameters in BGN^flox^ and BGN^ECKO^ mice following endothelial TXNDC5 overexpression (n=6). Data represent the mean±SD. Comparisons of parameters were performed with the unpaired 2-tailed Student *t*-test (**A-C**) or 2-way ANOVA (**D-F** and **H-K**), followed by Tukey honestly significant difference test for multiple comparisons. Nor indicates normoxia; Hyp, hypoxia; SuHx, and Sugen5416/hypoxia.

## References

1. Johnson S, Sommer N, Cox-Flaherty K, Weissmann N, Ventetuolo CE, Maron BA. Pulmonary Hypertension: A Contemporary Review. Am J Respir Crit Care Med. 2023;208:528–548.

2. Evans CE, Cober ND, Dai Z, Stewart DJ, Zhao Y-Y. Endothelial cells in the pathogenesis of pulmonary arterial hypertension. Eur Respir J. 2021;58:2003957.

3. Ma B, Cao Y, Qin J, Chen Z, Hu G, Li Q. Pulmonary artery smooth muscle cell phenotypic switching: A key event in the early stage of pulmonary artery hypertension. Drug Discov Today. 2023;28:103559.

4. Mandras SA, Mehta HS, Vaidya A. Pulmonary Hypertension: A Brief Guide for Clinicians. Mayo Clin Proc. 2020;95:1978–1988.

5. Olsson KM, Corte TJ, Kamp JC, Montani D, Nathan SD, Neubert L, Price LC, Kiely DG. Pulmonary hypertension associated with lung disease: new insights into pathomechanisms, diagnosis, and management. Lancet Respir Med. 2023;11:820–835.

6. Kanemura S, Matsusaki M, Inaba K, Okumura M. PDI Family Members as Guides for Client Folding and Assembly. Int J Mol Sci. 2020;21:9351.

7. Kuramochi T, Yamashita Y, Arai K, Kanemura S, Muraoka T, Okumura M. Boosting the enzymatic activity of CxxC motif-containing PDI family members. Chem Commun (Camb). 2024;60:6134–6137.

8. Hung C-T, Tsai Y-W, Wu Y-S, Yeh C-F, Yang K-C. The novel role of ER protein TXNDC5 in the pathogenesis of organ fibrosis: mechanistic insights and therapeutic implications. J Biomed Sci. 2022;29:63.

9. Jin Y, Sharma A, Bai S, Davis C, Liu H, Hopkins D, Barriga K, Rewers M, She J-X. Risk of type 1 diabetes progression in islet autoantibody-positive children can be further stratified using expression patterns of multiple genes implicated in peripheral blood lymphocyte activation and function. Diabetes. 2014;63:2506–2515.

10. Wang L, Dong H, Song G, Zhang R, Pan J, Han J. TXNDC5 synergizes with HSC70 to exacerbate the inflammatory phenotype of synovial fibroblasts in rheumatoid arthritis through NF-κB signaling. Cell Mol Immunol. 2018;15:685–696.

11. Wang X, Li H, Chang X. The role and mechanism of TXNDC5 in diseases. Eur J Med Res. 2022;27:145.

12. Yeh C-F, Cheng S-H, Lin Y-S, Shentu T-P, Huang R-T, Zhu J, Chen Y-T, Kumar S, Lin M-S, Kao H-L, et al. Targeting mechanosensitive endothelial TXNDC5 to stabilize eNOS and reduce atherosclerosis in vivo. Sci Adv. 2022;8:eabl8096.

13. Wu S, Luo X, Chen Y, Wang Z, Liu X, Sun N, Zhao J, Luo W, Zhang J, Tong X, et al. Sodium-glucose cotransporter 2 inhibitors attenuate vascular calcification by suppressing endoplasmic reticulum protein thioredoxin domain containing 5 dependent osteogenic reprogramming. Redox Biol. 2024;73:103183.

14. Fu Y, Zhou Y, Wang K, Li Z, Kong W. Extracellular Matrix Interactome in Modulating Vascular Homeostasis and Remodeling. Circ Res. 2024;134:931–949.

15. Thenappan T, Chan SY, Weir EK. Role of extracellular matrix in the pathogenesis of pulmonary arterial hypertension. Am J Physiol Heart Circ Physiol. 2018;315:H1322–H1331.

16. Nie X, Shen C, Tan J, Wu Z, Wang W, Chen Y, Dai Y, Yang X, Ye S, Chen J, et al. Periostin: A Potential Therapeutic Target For Pulmonary Hypertension? Circ Res. 2020;127:1138–1152.

17. Bian J-S, Chen J, Zhang J, Tan J, Chen Y, Yang X, Li Y, Deng L, Chen R, Nie X. ErbB3 Governs Endothelial Dysfunction in Hypoxia-Induced Pulmonary Hypertension. Circulation. 2024;150:1533–1553.

18. Robinson RM, Reyes L, Duncan RM, Bian H, Reitz AB, Manevich Y, McClure JJ, Champion MM, Chou CJ, Sharik ME, et al. Inhibitors of the protein disulfide isomerase family for the treatment of multiple myeloma. Leukemia. 2019;33:1011–1022.

19. Semenza GL. Hypoxia-inducible factors: roles in cardiovascular disease progression, prevention, and treatment. Cardiovasc Res. 2023;119:371–380.

20. Jiang Q, Braun DA, Clauser KR, Ramesh V, Shirole NH, Duke-Cohan JE, Nabilsi N, Kramer NJ, Forman C, Lippincott IE, et al. HIF regulates multiple translated endogenous retroviruses: Implications for cancer immunotherapy. Cell. 2025;188:1807–1827.e34.

21. Sullivan DC, Huminiecki L, Moore JW, Boyle JJ, Poulsom R, Creamer D, Barker J, Bicknell R. EndoPDI, a novel protein-disulfide isomerase-like protein that is preferentially expressed in endothelial cells acts as a stress survival factor. J Biol Chem. 2003;278:47079–47088.

22. Jandl K, Radic N, Zeder K, Kovacs G, Kwapiszewska G. Pulmonary vascular fibrosis in pulmonary hypertension - The role of the extracellular matrix as a therapeutic target. Pharmacol Ther. 2023;247:108438.

23. Sun X-J, Ma W-Q, Zhu Y, Liu N-F. POSTN promotes diabetic vascular calcification by interfering with autophagic flux. Cell Signal. 2021;83:109983.

24. Nikoloudaki G, Snider P, Simmons O, Conway SJ, Hamilton DW. Periostin and matrix stiffness combine to regulate myofibroblast differentiation and fibronectin synthesis during palatal healing. Matrix Biol. 2020;94:31–56.

25. Yu M, He X, Song X, Gao J, Pan J, Zhou T, Wang Q, Zhu W, Ma H, Zeng H, et al. Biglycan promotes hepatic fibrosis through activating heat shock protein 47. Liver Int. 2023;43:500–512.

26. Appunni S, Rubens M, Ramamoorthy V, Anand V, Khandelwal M, Sharma A. Biglycan: an emerging small leucine-rich proteoglycan (SLRP) marker and its clinicopathological significance. Mol Cell Biochem. 2021;476:3935–3950.

27. Prchal JT, Gordeuk VR. HIF-2 inhibitor, erythrocytosis, and pulmonary hypertension. Blood. 2021;137:2424–2425.

28. Shimoda LA. What’s HIF Got to Do with It? HIF-2 Inhibition and Pulmonary Hypertension. Am J Respir Crit Care Med. 2018;198:1363–1365.

29. Dai Z, Li M, Wharton J, Zhu MM, Zhao Y-Y. Prolyl-4 Hydroxylase 2 (PHD2) Deficiency in Endothelial Cells and Hematopoietic Cells Induces Obliterative Vascular Remodeling and Severe Pulmonary Arterial Hypertension in Mice and Humans Through Hypoxia-Inducible Factor-2α. Circulation. 2016;133:2447–2458.

30. Tang H, Babicheva A, McDermott KM, Gu Y, Ayon RJ, Song S, Wang Z, Gupta A, Zhou T, Sun X, et al. Endothelial HIF-2α contributes to severe pulmonary hypertension due to endothelial-to-mesenchymal transition. Am J Physiol Lung Cell Mol Physiol. 2018;314:L256–L275.

