## Supplemental Material 1 for "TXNDC5 Governs Extracellular Matrix Homeostasis in Pulmonary Hypertension"

### Expanded Methods and Materials

#### Study approval

All human samples were obtained from the Shenzhen People's Hospital (China) and Wuxi People's Hospital (China) with the informed consent of the patients, which was approved by the Ethics Committee for the Use of Human Subjects following The Code of Ethics of the Helsinki Declaration of the World Medical Association. All animal studies followed the guidelines of the Committee on Animal Research and Ethics of the Southern University of Science and Technology (China).

#### Clinical sample collection

Lung tissue and serum samples were obtained from patients with preoperative hypoxic pulmonary hypertension and matched healthy controls, all of whom were part of the lung transplantation cohort at Shenzhen People's Hospital and Wuxi People's Hospital. The healthy controls were individuals who were not suitable candidates for transplantation. Detailed clinical information for the samples can be found in **Table S1**.

#### Animal experiments design

Both male and female mice (C57BL/6 background) were used to perform the study. All animals were obtained from Cyagen (Jiangsu, China) and kept in a specific pathogen-free environment and a 12-hour light/dark cycle at room temperature. To generate homozygous TXNDC5 knockout mice ($TXNDC5^{-/-}$), the TXNDC5 knockout heterozygous mice ($TXNDC5^{+/-}$) were crossed with $TXNDC5^{+/-}$ to obtain $TXNDC5^{-/-}$ mice and their littermate control wild-type (WT) mice. Since BGN is located on the X chromosome, male mice carry a single floxed allele ($BGN^{f/0}$) and female mice carry two floxed alleles ($BGN^{f/f}$), and these mice are collectively referred to as $BGN^{flox}$ mice. To generate endothelial cell-specific TXNDC5 or BGN knockout mice ($TXNDC5^{ECKO}$ or $BGN^{ECKO}$), floxed TXNDC5 homozygous mice ($TXNDC5^{f/f}$) or floxed BGN mice ($BGN^{flox}$) were crossed with $Cdh5^{CreERT2}$ (Cre) mice. Mice were treated with tamoxifen (75 mg/kg, MedChemExpress, USA) via intraperitoneal injection once daily for 5 days.

For the SU5416/hypoxia (SuHx)-induced mice model, 8-12-week-old mice were injected subcutaneously with SU5416 (20 mg/kg, Sigma-Aldrich, USA) once per week for 3 weeks and exposed to hypoxia (10% $O_2$) in the ventilation chamber for 4 weeks. Mice in the control group were kept under normoxia for the same duration.

For *in vivo* overexpression assays, the pulmonary endothelium-specific serotype adeno-associated virus (AAV-LungX, manufactured by Obio Technology Co., Ltd., Shanghai, China) was used to deliver the Cdh5-Txndc5-3×FLAG-P2A-GdGreen-WPRE plasmid to achieve TXNDC5 overexpression. A total of $5\times10^{11}$ genome equivalent vectors were administered via tail vein injection once at the beginning of the 2 weeks before followed by SuHx exposure. For *in vivo* HIF-2α overexpression, as described by previous studies[1], two proline residues (Pro405 and Pro531) were substituted with alanine to prevent degradation, allowing stable HIF-2α expression under normoxic conditions. Then, Cdh5-HIF-2α (dPA)-3×FLAG-P2A-GdGreen-WPRE plasmid was delivered by adeno-associated virus to achieve endothelial-specific overexpression of stabilized HIF-2α. Control mice were injected with AAV carrying the empty vector (AAV-NC) at the same dose.

For the SuHx-induced rat model, male and female Sprague Dawley (SD) rats were injected subcutaneously with SU5416 (20 mg/kg) once, exposed to hypoxia (10% $O_2$) for 3 weeks, and then housed in normoxia for 2 weeks. For pharmacological inhibition of TXNDC5, after being exposed to SuHx for 3 weeks, rats were intraperitoneally injected once daily with E64FC26 (dissolved in saline containing 0.5% DMSO, 2 mg/kg) for 2 weeks. Control rats received vehicle injections instead under normoxia for the same duration. For TXNDC5-targeted gene therapy in rats, a self-complementary adeno-associated virus serotype LungX (scAAV-LungX, Obio Technology Co., Ltd., Shanghai, China) carrying a short hairpin RNA against TXNDC5 [Cdh5-shRNA(Txndc5)-CMV-EGFP-tWPA] or a non-targeting control shRNA was administered via a single intravenous injection through the tail vein at a dose of $5\times10^{12}$ genome equivalent after 2 weeks of exposure to SuHx. The rat-specific TXNDC5 shRNA utilized in this construct consisted of the sequence 5-TTTCTCAAGGCAACCACTTTA-3’.

**Echocardiography**

Transthoracic echocardiography was conducted using a VisualSonics Vevo 1100 ultrasound system (Toronto, Canada), equipped with an MS-400 probe for mice or an MS-250 for rats. Animals were anesthetized by continuous inhalation of isoflurane, and then fur was removed over their chest area. Pulmonary artery acceleration time (PAT) and pulmonary artery ejection time (PET) were measured using B-mode. Tricuspid annular plane systolic excursion (TAPSE), right ventricular fractional area change (RVFAC), and right ventricular free wall thickness (RVFWT) were assessed using M-mode echocardiography.

**Right ventricular systolic pressure (RVSP) measurement**

For RVSP measurement, animals were continuously anesthetized with inhalation of isoflurane, and a Millar Mikro-Tip pressure catheter (Millar Inc., Texas, USA) was surgically inserted into the right jugular vein and advanced into the right ventricle. Data was recorded using Power Lab monitoring equipment (AD Instruments, Germany).

**Pulmonary angiogram**

The mouse pulmonary angiogram was performed as previously described[2]. Chests were opened to expose the heart and lungs. Phosphate-buffered saline (containing heparin) was injected into the pulmonary artery via the RV, and the residual blood in the lungs was removed. Followed by perfusion with Microfil Injection Compounds (MV-122, Microfil, USA). Then, the lungs underwent gradient dehydration with ethanol, were subsequently immersed in methyl salicylate overnight, imaged by a stereomicroscope, and analyzed by using Image J.

**Histological analysis of lung tissues**

Tissue specimens were fixed in 4% formaldehyde, dehydrated in graded ethanol, transparentized in xylene, and embedded in paraffin. Lung tissue sections were stained with hematoxylin and eosin (H&E) to visualize general tissue morphology and cellular architecture, followed by scanning with the Aperio VERSA 8 system (Leica, USA). Pulmonary arterial medial wall thickness was measured using ImageJ software.

For immunohistochemical (IHC) analysis, α-SMA (Cell Signaling Technology, 19245S) was used to visualize smooth muscle cells in pulmonary arteries. The α-SMA-positive area within the vessel media was quantified using ImageJ software, and the proportion of α-SMA-positive area relative to the total medial area was calculated. Pulmonary arteries were classified as fully muscularized (>75%), partially muscularized (25-75% α-SMA-positive area), or non-muscularized (<25%). At least 20 vessels (25-75 mm diameter) per animal were analyzed, and the percentages of vessels in each category were calculated to evaluate pulmonary arterial muscularization.

For immunofluorescence staining (IF) analysis, tissue sections were first deparaffinized and rehydrated. Endogenous peroxidase activity was subsequently quenched, followed by blocking with 2% BSA for 1 hour at room temperature. After blocking, the sections were incubated with primary antibodies at 4°C overnight (TXNDC5, Proteintech, 19834-1-AP, 1:200; α-SMA, Cell Signaling Technology, 19245S, 1:200; CD31, Dianova, DIA-310, 1:100; BGN, Proteintech, 16409-1-AP, 1:100; POSTN, Novus, NBP1-30042, 1:100**;** COL1A1, A24112, 1:500, ABclonal; HIF-2α, NB100-122, 1:200). Then, the sections were incubated with the corresponding Alexa Fluor-conjugated secondary antibodies for 1 hour at room temperature in the dark. For multiplex immunofluorescence (mIF) staining analysis, the sections were stained using an mIF kit (AFIHC025, Hunan Aifang Biological Technology, China) based on the tyramide signal amplification technology according to the manufacturer's instructions. All sections were observed using an LSM980 laser confocal microscope (Zeiss, Germany). The mean fluorescence intensity (MFI) was measured using ImageJ software.

**Right ventricular (RV) hypertrophy and fibrosis evaluation**

For RV hypertrophy evaluation, the whole heart was isolated, and the right ventricular (RV) was carefully dissected from the left ventricle and septum (LV+S). The Fulton index was calculated with the weight ratio of RV/(LV+S) to evaluate RV hypertrophy. Meanwhile, heart sections were stained by H&E and then scanned by Aperio VERSA

8 (Leica, USA). The thickness of the RV wall was measured with ImageJ software. For wheat germ agglutinin (WGA) staining of mouse right ventricles, specimens were fixed with 4% formaldehyde overnight and embedded in paraffin, sectioned at 5 μm. After deparaffinizing, rehydrating, and blocking, sections were stained with Alexa Fluor 488-labeled WGA (W11261, Sigma-Aldrich, 1:200 dilution) for 2 h at room temperature. Nuclei were stained with DAPI for 10 min. All sections were observed using an LSM980 laser confocal microscope (Zeiss, Germany). The cardiomyocyte cross-sectional area (CSA) was analyzed and quantified by ImageJ.

**E64FC26 toxicity assessment**

Rats were intraperitoneally injected once daily with E64FC26 (2 mg/kg) for 5 weeks under normoxic conditions. Body weight was measured every other day during the treatment period using a calibrated digital scale. The liver, kidney, and spleen tissues were fixed and then subjected to dehydration, clearing, and paraffin embedding. Tissue sections were stained with H&E and scanned by Aperio VERSA 8.

***In vivo* imaging**

Major organs (heart, liver, spleen, lung, and kidney) of animals were harvested, rinsed in PBS, and placed in the imaging chamber. Fluorescence signals were acquired using GFP excitation/emission filters with standardized exposure times. Images were captured and analyzed using an IVIS Spectrum system (Caliper Life Sciences, USA).

**Lung endothelial cell isolation**

Mouse lung endothelial cell isolation was performed and modified as previously described[3]. In brief, lung tissues were harvested, minced, and digested in 2 mg/ml Collagenase A (Sigma-Aldrich) for 40 min at 37°C. Cell precipitates were then incubated with CD31 (Dianova, DIA-310)-conjugated Dynabeads (11035, Sigma-Aldrich) for 15 min at room temperature. After washing, beads were suspended and then seeded in gelatin-coated 6-well plates till cells reached confluence. For rat lung endothelial cell isolation, lung tissues were harvested, minced, and digested in 2 mg/ml

Collagenase A for 1 h at 37°C. Cell suspensions were incubated with CD31 (Servicebio, GB11063-2)-conjugated Dynabeads (11203, Sigma-Aldrich), washed, and seeded in gelatin-coated 6-well plates until confluence.

**Cell lines and cell culture**

Human pulmonary endothelial cells (PAECs) were obtained from Meisen Chinese Tissue Culture Collections (Zhejiang, China). Cells in passages 3-8 were used in all experiments. PAECs and primary lung ECs were cultured in endothelial cell medium (1001, ScienCell, USA). HEK293T cells were obtained from ATCC and maintained in DMEM (11965092, Gibco, USA). All cells were maintained at 37°C in 95% air and 5% $CO_2$. For hypoxia exposure, cells were incubated in a Variable Oxygen Control $CO_2$ incubator (Thermo Fisher Scientific, USA) with an oxygen concentration of 1%.

**Western blotting analysis and immunoprecipitation**

Proteins from cells or tissues were extracted by RIPA lysis buffer (P0013B, Beyotime, China) supplemented with 1mM PMSF (ST507, Beyotime). Lysates were separated by SDS-PAGE and transferred to polyvinylidene fluoride membranes. Membranes were immunoblotted with primary antibodies (TXNDC5, Proteintech, 19834-1-AP, 1:5000; BGN (Rabbit), Proteintech, 16409-1-AP, 1:1000; BGN (Mouse), Santa Cruz, sc-100857, 1:500; Vinculin, Proteintech, 66305-1-Ig, 1:5000; β-actin, AC004, ABclonal, 1:10000; HIF-1α, Proteintech, 66730-1-Ig, 1:2000; HIF-2α, HUABIO, ET7107-32, 1:1000; HA, Proteintech, 51064-2-AP, 1:5000; FLAG, Abmart, M20008, 1:5000; POSTN, Novus, NBP1-30042, 1:1000; LOX, ABclonal, A11504, 1:1000; TGM2, ABclonal, A0981, 1:1000; FN1, F3648, 1:5000) overnight at 4 °C. For immunoprecipitation, specific antibodies (TXNDC5, Proteintech, 19834-1-AP; HA, Proteintech, 51064-2-AP; and IgG (Rabbit), 2729S, Cell Signaling Technology) were incubated with Protein A/G Magnetic Beads (MedChemExpress, USA) for 2 h at 4°C, followed by incubation with lysates for 2 h at 4°C. Immunoprecipitated products were separated by SDS-PAGE and immunoblotted with the antibodies as indicated.

**Chromatin immunoprecipitation assay (ChIP) and quantitative polymerase chain reaction (qPCR)**

ChIP was performed according to the manufacturer's instructions (P2078, Beyotime, China). In brief, cells were crosslinked with 1% formaldehyde for 10 minutes, followed by quenching with glycine. Chromatin was extracted and fragmented by sonication to an average size of 200-500 bp. The lysates were incubated with specific antibodies (HIF-2α, Cell Signaling Technology, 87179S, and IgG (Rabbit), 2729S, Cell Signaling Technology) overnight at 4°C. Immunoprecipitated complexes were captured with Protein A/G beads, washed, and eluted. Immunoprecipitated DNA fragments were amplified by qPCR and visualized by agarose gel electrophoresis. For qPCR, the total RNA of samples was extracted using RNAiso Plus (9108, Takara, Japan). The mRNA level was measured by using SYBR Green (Yeason, China) with β-actin as an internal control. Primer sequences used for qPCR and ChIP-qPCR are in **Table S2**.

**Ethynyl-2′-deoxyuridine (EdU) staining**

As follows by the manufacturer's instructions (C0075S, Beyotime, China), cells were incubated with 10 μM EdU for 2 hours at 37°C. After incubation, cells were fixed with 4% paraformaldehyde, permeabilized with 0.5% Triton X-100, and then incubated with the EdU detection solution. Nuclei were counterstained with Hoechst 33342, and images were captured using a laser confocal microscope.

**Cell cycle analysis**

Cell cycle analysis was performed by using the Cell Cycle and Apoptosis Analysis Kit (C1052, Beyotime, China). Cells were harvested and fixed with 70% ethanol overnight at 4°C. After fixation, cells were washed and incubated with propidium iodide (PI) staining solution, and the cells were incubated for 30 min at room temperature in the dark. Cell cycle distribution was analyzed by flow cytometry using the FACSCanto SORP (BD, USA), and data were analyzed using FlowJo software.

**Cellular immunofluorescence staining**

Cells were fixed, permeabilized, blocked, and then incubated with primary antibodies (TXNDC5, Proteintech, 19834-1-AP, 1:200; BGN, Abcam, ab58562, 1:100; HIF-2α, Proteintech, 26422-1-AP, 1:100; Phalloidin, Invitrogen, A12379, 1:2000; and Paxillin, Proteintech, 22172-1-AP, 1:500) overnight at 4°C. Nuclei were stained with DAPI, and the cells were imaged using a laser confocal microscope.

**Cell migration assay**

Cells were plated in 6-well plates and grown to confluence, and the culture medium was replaced with serum-free medium. A sterile pipette tip was used to create a straight-line scratch in the monolayer. After scratching, cells were incubated at 37°C, and images were captured at 0, 12, 24, and 48 h to monitor wound closure. The average cell migration distance was measured, and the relative migration ratio was quantified using ImageJ.

**Cell transfection and virus infection**

For small interfering RNA (siRNA) transfection, 20 nM TXNDC5, HIF-2α, or control siRNA (RiboBio, China) was diluted in Opti-MEM medium (31985070, Gibco, USA) and transfected into cells with the use of Lipofectamine RNAiMAX (13778150, Invitrogen, USA). For lentivirus (LV) delivered overexpression plasmids transfection, cells were incubated with LV-HIF-2α (dPA) or LV-TXNDC5 or LV-TXNDC5 (Mut) or control LV (LV-NC) (OBIO, China) at a multiplicity of infection (MOI) of 20, along with 10 μg/mL polybrene (MedChemExpress, USA) to enhance infection efficiency. Infected cells were incubated for 48-72 h, and gene overexpression was confirmed by western blot or qPCR. For immunoprecipitation, FLAG-BGN and HA-TXNDC5 plasmids were transfected into HEK293T by using the Lipofectamine 3000 (L3000015, Invitrogen, USA).

TXNDC5 Mut sequences with cysteine (C)-to-alanine (A) substitutions in all three thioredoxin domains. TXNDC5 WT: N'---CGHC---CGHC---CGHC---C'; TXNDC5 Mut: N'---AGHA---AGHA---AGHA---C'.

**Cycloheximide (CHX) chase assay**

Cells were seeded in six-well plates and treated with cycloheximide (CHX, 50 μg/mL) to inhibit protein synthesis. They were harvested at different time points after CHX treatment. Protein levels were analyzed by Western blot to assess protein degradation kinetics.

**Dot bolt assays for secreted proteins**

The same volume of cell supernatants in different groups was concentrated by using the Amicon Ultra Centrifugal Filters (Millipore, USA). Then, 5 μL samples were dotted onto the nitrocellulose membrane, dried, ultraviolet crosslinked, blocked, and incubated with specific primary antibodies. The membrane was visualized with enhanced chemiluminescence reagents.

**Enzyme-linked immunosorbent assay (ELISA)**

The levels of BGN in the serum from PH patients were measured with the Human Biglycan ELISA Kit (JONLNBiO, China) according to the manufacturer's instructions. Briefly, 100 μL samples were incubated for 1 h at 37°C, and the biotinylated antibody solution was added to incubate for 1 h. After washing, the enzyme conjugate working solution was then added for 30 min at 37°C, followed by incubation with TMB substrate for 15 min in the dark. The absorbance at 450 nm was measured using a microplate reader (BioTek, USA).

**Protein-protein interaction (PPI)**

For TXNDC5-targeted extracellular matrix (ECM) PPI network construction, differentially expressed ECM proteins were identified at a 1.5-fold change and $p$<0.05 (Student $t$-test). Differentially expressed ECM proteins were visualized by a heatmap using the R language. Then, TXNDC5-targeted ECM proteins were classified into different functional subgroups, including "matrix modulators and adhesion", "metallopeptidase activity", "basement membrane", "proteoglycans and glycoproteins", "growth factors and signaling regulators", and "Angiogenesis" according to the Gene

Ontology database (https://www.geneontology.org/) and Uniprot database (https://www.uniprot.org/). Detailed information was available in **Supplemental Material 4**. For ECs-function-ECM PPI network construction, we obtained a dataset of ECs function-related genes (including endothelium development, regulation of endothelial cell differentiation, regulation of endothelial cell chemotaxis, regulation of endothelial cell migration, regulation of endothelial cell apoptotic process, endothelial cell migration, endothelial cell differentiation, endothelial cell proliferation, endothelial cell activation, and endothelial cell development) and a dataset of genes encoding ECM proteins from Gene Ontology database. The above datasets were then mapped onto the PPI network by STRING (https://cn.string-db.org/) to identify interactions and visualized by Cytoscape software. Then, we reconstructed the genes with degree values greater than the average in each network to gradually get a new PPI network, and finally obtained the core network composed of the final hub genes. Detailed information was available in **Supplemental Material 5**.

**Dual-luciferase reporter assay**

HIF-2α expression plasmid and TXNDC5 promoter reporter plasmids (with or without the indicated mutations) were transfected into HEK293T cells. Luciferase activity was measured according to the instructions of the Dual-Luciferase Reporter Gene Assay Kit (RG027, Beyotime, China). Luminescence was detected by GloMax Navigator (GM2010, Promega, USA).

**Fluorescence resonance energy transfer (FRET)-based protein folding assay**

Protein folding assays were performed based on FRET as described previously[4]. Briefly, A plasmid encoding human BGN fused with CFP at the N-terminal and YFP at the C-terminal was constructed (CFP-BGN-YFP). CFP-BGN-YFP was transduced into cells to determine the protein folding efficiency of BGN through the standard acceptor photobleaching FRET protocol. An excitation wavelength of 458 nm and an emission wavelength of 465-515 nm were used for CFP, whereas an excitation wavelength of 514 nm and an emission wavelength of 525-555 nm were used for YFP. The FRET signals were collected by using an LSM980 laser confocal microscope (Zeiss, Germany) and quantified by using the Image J plugin AccPbFRET.

**Molecular docking**

The 3D structure of BGN and TXNDC5 was modeled by AlphaFold2 (https://alphafold.com/) and subjected to ZDOCK 3.0.2 for protein-protein docking. The protein-protein interaction was analyzed by the Protein-Ligand Interaction Profiler. The best docking conformations were selected based on the highest score and visualized by PyMOL software.

**4D label-free proteomics**

Proteins were extracted from lung tissues using SDT buffer (4% SDS, 100 mM Tris-HCl, 1 mM DTT, pH 7.6) and quantified with the BCA assay. Digestion was performed according to the filter-aided sample preparation (FASP) method, where proteins were treated with trypsin, and the resulting peptides were desalted using C18 cartridges (bed I.D. 7 mm, volume 3 ml, Sigma-Aldrich). Proteins were separated via SDS-PAGE and visualized by Coomassie Blue staining. For LC-MS/MS analysis, a Q Exactive mass spectrometer was coupled with Easy nLC (Thermo Scientific). Peptides were loaded onto a C18 trap column and separated on a C18 analytical column (10 cm, 75 μm inner diameter, 3 μm resin, Thermo Scientific). MS data were collected in positive ion mode with data-dependent acquisition for HCD fragmentation, using a resolution of 70,000 at m/z 200 and dynamic exclusion of precursor ions. Raw data were processed and analyzed using MaxQuant software for protein identification and quantification. Hierarchical clustering was performed with Cluster 3.0 and Java Treeview (http://jtreeview.sourceforge.net) using Euclidean distance and average linkage clustering algorithms. Motif analysis was conducted with MeMe (https://meme-suite.org/meme/). Detailed information was available in **Supplemental Material 2**.

**RNA-sequencing and data analysis**

PAECs were transfected with TXNDC5 siRNA or control siRNA under hypoxic conditions, and total RNA was extracted with RNAiso Plus. RNA samples were detected based on the A260/A280 absorbance ratio with a Nanodrop ND-2000 system (Thermo Scientific), and the RIN of RNA was determined by an Agilent Bioanalyzer 4150 system (Agilent Technologies, CA, USA). Paired-end libraries were prepared using an mRNA-seq Lib Prep Kit (ABclonal, China) following the manufacturer's instructions. The library preparations were sequenced on an Illumina Novaseq 6000, and 150 bp paired-end reads were generated. Raw reads of fastq format were first processed through in-house Perl scripts, and then clean reads were separately aligned to the reference genome with orientation mode using HISAT2 software to obtain mapped reads. The FPKM of each gene was calculated based on the length of the gene and the read count mapped to the gene. Genes were identified as significantly differentially expressed at 1.5-fold change and $p<0.05$ (Student *t*-test). Enrichment analysis of differentially expressed genes was performed using the DAVID database (https://davidbioinformatics.nih.gov/) and visualized by a bubble chart using the R language. Detailed information was available in **Supplemental Material 3**.

**Statistical Analysis**

Data were analyzed with GraphPad Prism 8.0 software and were expressed as the mean±Standard Deviation (mean±SD). For comparisons between two groups with a normal distribution, an unpaired 2-tailed Student *t*-test was performed. When the data did not follow a normal distribution, the non-parametric Mann-Whitney *U* test was applied. For data involving more than two groups, one-way ANOVA was used, followed by Tukey's honestly significant difference test for multiple comparisons. For two variables, two-way ANOVA with Tukey's honestly significant difference test for multiple comparisons was applied. Statistical significance was defined as $P<0.05$.

### References

1. Jiang Q, Braun DA, Clauser KR, Ramesh V, Shirole NH, Duke-Cohan JE, et al. HIF regulates multiple translated endogenous retroviruses: Implications for cancer immunotherapy. Cell 2025;188:1807-1827.e34.

doi:10.1016/j.cell.2025.01.046.

2. Xiong M, Jain PP, Chen J, Babicheva A, Zhao T, Alotaibi M, et al. Mouse model of experimental pulmonary hypertension: Lung angiogram and right heart catheterization. Pulm Circ 2021;11:20458940211041512. doi:10.1177/20458940211041512.

3. Wang J, Niu N, Xu S, Jin ZG. A simple protocol for isolating mouse lung endothelial cells. Sci Rep 2019;9:1458. doi:10.1038/s41598-018-37130-4.

4. Shih Y-C, Chen C-L, Zhang Y, Mellor RL, Kanter EM, Fang Y, et al. Endoplasmic Reticulum Protein TXNDC5 Augments Myocardial Fibrosis by Facilitating Extracellular Matrix Protein Folding and Redox-Sensitive Cardiac Fibroblast Activation. Circ Res 2018;122:1052–1068. doi:10.1161/CIRCRESAHA.117.312130.

**Table S1. Clinical characteristics of PH patients and donors**

| Sample ID | Assays | Age (year) | Sex | Race/ Ethnicity | Diagnosis/ Cause of death | sPAP (mmHg) |
|---|---|---|---|---|---|---|
| PH-1 | IF, WB (lung tissue) | 29 | F | Asian | COPD-associated PH | 79 |
| PH-2 | WB (lung tissue) | 56 | M | Asian | Pulmonary fibrosis-associated PH | 95 |
| PH-3 | IF, WB (lung tissue) | 63 | M | Asian | COPD-associated PH | 68 |
| PH-4 | IF, WB (lung tissue) | 47 | M | Asian | COPD-associated PH | 133 |
| PH-5 | IF, WB (lung tissue) | 51 | M | Asian | Interstitial lung disease-associated PH | 110 |
| PH-6 | IF, WB (lung tissue) | 32 | F | Asian | Interstitial lung disease-associated PH | 54 |
| PH-7 | WB (lung tissue) | 58 | F | Asian | Silicosis PH | 87 |
| PH-8 | IF (lung tissue) | 60 | M | Asian | COPD-associated PH | 76 |
| PH-9 | IF (lung tissue) | 48 | F | Asian | COPD-associated PH | 68 |
| PH-10 | IF (lung tissue) | 72 | F | Asian | Interstitial lung disease-associated PH | 105 |
| PH-11 | IF (lung | 55 | M | Asian | Interstitial | 59 |

| | | | | | | |
|---|---|---|---|---|---|---|
| | tissue) | | | | lung disease-associated PH | |
| PH-12 | IF (lung tissue) | 61 | M | Asian | Interstitial lung disease-associated PH | 77 |
| PH-13 | ELISA (Serum) | 34 | F | Asian | Pulmonary fibrosis-associated PH | 39 |
| PH-14 | ELISA (Serum) | 72 | M | Asian | Pulmonary fibrosis-associated PH | 44 |
| PH-15 | ELISA (Serum) | 42 | M | Asian | Interstitial pneumonia-associated PH | 42 |
| PH-16 | ELISA (Serum) | 57 | F | Asian | Pulmonary fibrosis-associated PH | 38 |
| PH-17 | ELISA (Serum) | 54 | M | Asian | Pulmonary fibrosis-associated PH | 43 |
| PH-18 | ELISA (Serum) | 58 | M | Asian | Pulmonary fibrosis-associated PH | 47 |
| PH-19 | ELISA (Serum) | 57 | M | Asian | Bronchiectasis-associated PH | 41 |
| PH-20 | ELISA (Serum) | 55 | M | Asian | Interstitial lung disease-associated PH | 52 |

| | | | | | | |
|---|---|---|---|---|---|---|
| PH-21 | ELISA (Serum) | 38 | F | Asian | Pulmonary fibrosis-associated PH | 78 |
| PH-22 | ELISA (Serum) | 68 | M | Asian | Pulmonary fibrosis-associated PH | 74 |
| PH-23 | ELISA (Serum) | 39 | M | Asian | Silicosis PH | 54 |
| PH-24 | ELISA (Serum) | 72 | M | Asian | Pulmonary fibrosis-associated PH | 44 |
| PH-25 | ELISA (Serum) | 43 | M | Asian | Interstitial lung disease-associated PH | 61 |
| PH-26 | ELISA (Serum) | 62 | M | Asian | Pulmonary fibrosis-associated PH | 54 |
| PH-27 | ELISA (Serum) | 49 | F | Asian | Interstitial lung disease-associated PH | 71 |
| PH-28 | ELISA (Serum) | 52 | M | Asian | Pulmonary fibrosis-associated PH | 72 |
| PH-29 | ELISA (Serum) | 62 | M | Asian | Interstitial lung disease-associated PH | 80 |
| PH-30 | ELISA (Serum) | 32 | F | Asian | PVOD-associated PH | 105 |
| PH-31 | ELISA | 53 | M | Asian | Pulmonary | 82 |

| | | | | | | |
|---|---|---|---|---|---|---|
| | (Serum) | | | | fibrosis-associated PH | |
| PH-32 | ELISA (Serum) | 65 | M | Asian | COPD-associated PH | 58 |
| PH-33 | ELISA (Serum) | 60 | F | Asian | Interstitial lung disease-associated PH | 55 |
| PH-34 | ELISA (Serum) | 54 | M | Asian | Pulmonary fibrosis-associated PH | 57 |
| PH-35 | ELISA (Serum) | 41 | M | Asian | IPF-associated PH | 52 |
| PH-36 | ELISA (Serum) | 58 | M | Asian | Pulmonary fibrosis-associated PH | 81 |
| PH-37 | ELISA (Serum) | 60 | M | Asian | Interstitial lung disease-associated PH | 51 |
| PH-38 | ELISA (Serum) | 51 | M | Asian | Interstitial lung disease-associated PH | 30 |
| PH-39 | ELISA (Serum) | 64 | M | Asian | Pulmonary fibrosis-associated PH | 61 |
| Donor-1 | IF, WB (lung tissues) | 47 | M | Asian | N/A | N/A |
| Donor-2 | IF, WB (lung | 54 | F | Asian | N/A | N/A |

| | | | | | | |
|---|---|---|---|---|---|---|
| | tissues) | | | | | |
| Donor-3 | IF, WB (lung tissues) | 45 | M | Asian | N/A | N/A |
| Donor-4 | IF, WB (lung tissues) | 42 | M | Asian | N/A | N/A |
| Donor-5 | IF, WB (lung tissues) | 32 | M | Asian | N/A | N/A |
| Donor-6 | WB (lung tissues) | 39 | F | Asian | N/A | N/A |
| Donor-7 | WB (lung tissues) | 36 | F | Asian | N/A | N/A |
| Donor-8 | ELISA (Serum) | 42 | F | Asian | N/A | N/A |
| Donor-9 | ELISA (Serum) | 40 | M | Asian | N/A | N/A |
| Donor-10 | ELISA (Serum) | 28 | M | Asian | N/A | N/A |
| Donor-11 | ELISA (Serum) | 25 | F | Asian | N/A | N/A |
| Donor-12 | ELISA (Serum) | 58 | M | Asian | N/A | N/A |
| Donor-13 | ELISA (Serum) | 50 | M | Asian | N/A | N/A |
| Donor-14 | ELISA (Serum) | 61 | M | Asian | N/A | N/A |
| Donor-15 | ELISA | 71 | F | Asian | N/A | N/A |

| | | | | | | |
|---|---|---|---|---|---|---|
| | (Serum) | | | | | |
| Donor-16 | ELISA (Serum) | 38 | F | Asian | N/A | N/A |
| Donor-17 | ELISA (Serum) | 32 | M | Asian | N/A | N/A |
| Donor-18 | ELISA (Serum) | 46 | M | Asian | N/A | N/A |
| Donor-19 | ELISA (Serum) | 62 | M | Asian | N/A | N/A |
| Donor-20 | ELISA (Serum) | 55 | F | Asian | N/A | N/A |
| Donor-21 | ELISA (Serum) | 52 | F | Asian | N/A | N/A |
| Donor-22 | ELISA (Serum) | 29 | M | Asian | N/A | N/A |
| Donor-23 | ELISA (Serum) | 28 | F | Asian | N/A | N/A |
| Donor-24 | ELISA (Serum) | 37 | M | Asian | N/A | N/A |
| Donor-25 | ELISA (Serum) | 36 | M | Asian | N/A | N/A |
| Donor-26 | ELISA (Serum) | 61 | M | Asian | N/A | N/A |
| Donor-27 | ELISA (Serum) | 63 | F | Asian | N/A | N/A |
| Donor-28 | ELISA (Serum) | 58 | F | Asian | N/A | N/A |
| Donor-29 | ELISA (Serum) | 55 | M | Asian | N/A | N/A |

| | | | | | | |
|---|---|---|---|---|---|---|
| Donor-30 | ELISA (Serum) | 50 | M | Asian | N/A | N/A |
| Donor-31 | ELISA (Serum) | 40 | M | Asian | N/A | N/A |
| Donor-32 | ELISA (Serum) | 42 | F | Asian | N/A | N/A |
| Donor-33 | ELISA (Serum) | 60 | M | Asian | N/A | N/A |
| Donor-34 | ELISA (Serum) | 56 | F | Asian | N/A | N/A |

Definition of abbreviations: F, female; M, male; PH, pulmonary hypertension; COPD, chronic obstructive pulmonary disease; IPF, idiopathic pulmonary fibrosis; PVOD, pulmonary venoocclusive disease; sPAP, mean systolic pulmonary artery pressure; N/A, data not available.

**Table S2. Primer sequences used for qPCR and ChIP-qPCR**

| Sites | **Forward** | **Reverse** |
|---|---|---|
| TXNDC5 | CAGAGCCGGAAGTGGAACC | CCACGGAGCGAAGAACTTGAT |
| BGN | CAGTGGCTTTGAACCTGGAG | GGGAGGTCTTTGGGGATGC |
| POSTN | CTCATAGTCGTATCAGGGGTCG | ACACAGTCGTTTTCTGTCCAC |
| COL1A1 | GTGCGATGACGTGATCTGTGA | CGGTGGTTTCTTGGTCGGT |
| FN1 | CAGCCTTCCACTAAGGATTTGC | GCTCTAAACCCCTTCTCTCTGAG |
| VEGFA | AGGGCAGAATCATCACGAAGT | AGGGTCTCGATTGGATGGCA |
| EDN1 | AGAGTGTGTCTACTTCTGCCA | CTTCCAAGTCCATACGGAACAA |
| β-actin | CATGTACGTTGCTATCCAGGC | CTCCTTAATGTCACGCACGAT |
| Binding site 1 | AGAGCAACCCGGTGATTCTAA | AAAATGAATGACAGCACGCC |
| Binding site 2 | CAGTCCTGCTAGGTGCCTTC | TCTTGGCCAGAGCTCAACTG |
